# Epigenomic and transcriptomic prioritization of candidate obesity-risk regulatory GWAS SNPs

**DOI:** 10.1101/2021.10.29.466522

**Authors:** Xiao Zhang, Tianying Li, Hong-Mei Xiao, Kenneth C. Ehrlich, Hui Shen, Hong-Wen Deng, Melanie Ehrlich

**Affiliations:** Tulane Center for Biomedical Informatics and Genomics, Division of Biomedical Informatics and Genomics, Deming Department of Medicine, Tulane University, New Orleans, LA, USA; Center for System Biology, Data Sciences, and Reproductive Health, School of Basic Medical Science, Central South University, Changsha, Hunan, China; Tulane Cancer Center and Hayward Genetics Center, Tulane University, New Orleans, LA, USA

**Keywords:** GWAS, epigenetics, obesity, SNPs, lncRNA, RNA isoforms, enhancers, promoters, transcriptomics, subcutaneous adipose tissue

## Abstract

Concern about rising rates of obesity has prompted searches for its genetic risk determinants in genome-wide association studies (GWAS). Most genetic variants that contribute to the increased risk of a given trait are probably regulatory single nucleotide polymorphisms (SNPs). However, identifying plausible regulatory SNPs is difficult because of their varied locations relative to their target gene and linkage disequilibrium, which makes most GWAS-derived SNPs only proxies for many fewer functional SNPs. We developed a systematic approach to prioritizing GWAS-derived obesity SNPs using detailed epigenomic and transcriptomic analysis in adipose tissue vs. heterologous tissues. From 50 obesity-related GWAS and 121,064 expanded SNPs, we prioritized 47 potential causal regulatory SNPs (Tier-1 SNPs) for 14 gene loci. A detailed examination of seven of these genes revealed that four (*CABLES1, PC, PEMT,* and *FAM13A*) had Tier-1 SNPs that might regulate alternative use of transcription start sites resulting in different polypeptides being generated or different amounts of an intronic microRNA gene being expressed. *HOXA11* and long noncoding RNA gene *RP11-392O17.1* had Tier-1 SNPs in their 3’ or promoter region, respectively, and strong preferences for expression in subcutaneous vs. visceral adipose tissue. *ZBED3-AS1* had two intragenic Tier-1 SNPs, each of which might contribute to mediating obesity risk through modulating long-distance chromatin interactions. We conclude that prioritization of regulatory SNP candidates should focus on their surrounding epigenetic features in a trait-relevant tissue. Our approach not only revealed especially credible novel regulatory SNPs, but also helped evaluate previously highlighted obesity GWAS SNPs that were candidates for transcription regulation.

## Introduction

The rapidly rising rates of obesity are a major concern for public health and increased risk of disease, including type 2 diabetes (T2D), cardiovascular disease, osteoarthritis, certain kinds of cancer, and respiratory problems (1, 2). Adipose tissue expansion involves an increase in adipocytes and/or an expansion of the size of adipocytes. However, not only overall obesity but also the location of the fat depot with increased adipose tissue is of importance to health risk (2). The major types of fat depots are visceral adipose tissue (VAT), which is deposited around internal organs in the abdominal cavity and subcutaneous adipose tissue (SAT), which is beneath the skin. High levels of VAT are associated with metabolic abnormalities while high levels of SAT may be protective rather than harmful (3). Similarly, adipose deposition around the hips, which mostly involves SAT, is less likely to be a risk factor for various chronic diseases than are high levels of adipose at the waist, which have a large VAT component. Pre-menopausal women with their tendency to high SAT/VAT ratios are at lower risk of adipose-related disease (4). Nonetheless, expansion of SAT depots may also contribute to health either in its protective role or through its interactions with VAT depots (3, 5).

The clinical relevance of the location of fat depots in obesity led to more detailed measurements of obesity than just body mass index (BMI), namely, the waist-to-hip ratio (WHR) adjusted for BMI (WHR_adjBMI_), waist circumference adjusted for BMI (WC_adjBMI_), and hip circumference adjusted for BMI (HC_adjBMI_). These anthropomorphic measures are the most frequent ones used in genome-wide association studies of the genetic risk of obesity (obesity GWAS). Genetic risk clearly contributes to obesity, and genetic heritability for BMI has been estimated to be 30 – 40% (2), but this estimate depends on the study design (6).

Especially challenging in the analysis of most GWAS-derived genetic variants is assigning a biochemical function to them when they are not exonic and predicted to change the encoded protein’s activity or stability. For example, less than 10% of single nucleotide polymorphisms (SNPs, the most frequent genetic variants) significantly associated with obesity in GWAS are in coding regions (7), and only a fraction of these SNPs are at non-synonymous codons predicted to affect polypeptide structure. Most of the GWAS-derived SNPs that actually mitigate genetic risk for a given trait are probably in enhancers or, less frequently, promoters (8). The detection of regulatory SNPs is complicated by the variety of sizes and genomic locations for cell-type specific enhancers. Especially difficult is the finding that there are often tens to hundreds of SNPs tightly linked to an individual GWAS-reported SNP. Complex linkage disequilibrium (LD) between SNPs and causative mutations compounded by sampling errors in test statistics (9) obscures the assignment of a given SNP to a biological function.

To address the problems of identifying credible candidates for regulatory SNPs associated with obesity risk, we used a novel systematic approach dependent upon epigenomics and transcriptomics of not just a target tissue for obesity, adipose tissue, but also comparisons of adipose tissue to many other tissue types. This approach avoids the ambiguity introduced by LD, as seen, e. g., in assignments of expression quantitative trait loci (eQTLs), because epigenetic marks are independent of LD and directly related transcription regulation. While there has been more attention recently to epigenetics in studies of obesity and obesity risk (e.g., (1, 2, 10, 11)), our prioritization of highly plausible regulatory SNPs for obesity risk uses strict criteria for adipose-preferential epigenomics and transcriptomics as well as genetic criteria and predictions of allele-specific transcription factor (TF) binding. Although diverse non-adipose organ systems can contribute to obesity (e.g., brain, pancreas, liver, and bone), we focused on adipose tissue due to its direct relationship to obesity and the importance of adipose gene expression to obesity measurements (1, 2). From 50 obesity GWAS, we highlighted seven gene loci associated with 18 highly credible, mostly novel obesity-risk regulatory SNPs. These SNPs are candidates for modulating enhancers, promoters, or long-distance chromatin interactions affecting expression of protein-coding or long non-coding RNA (lncRNA) genes relevant to adipose biology.

## Results

### Candidate regulatory SNPs (Tier-1 SNPs) for obesity risk were prioritized from 50 obesity-related GWAS

To identify highly credible candidates for causative regulatory variants from obesity-related GWAS, index SNPs from 50 studies of BMI, WHR_adjBMI_, WC_adjBMI_, or HC_adjBMI_, (Supplementary Table S1) were expanded (LD threshold of *r*^2^ ≥ 0.8, European population, EUR), and the index or proxy SNPs (I/P SNPs) were prioritized using epigenomics, transcriptomics, and transcription factor binding site (TFBS) predictions (Figure 1). The prioritization depended on the assumptions that many of the regulatory SNPs affecting obesity risk should modulate transcription of genes preferentially expressed in adipose tissue, should directly overlap epigenetic regulatory features seen preferentially in adipose tissue, and should overlap a TFBS that binds to its TF selectively from one of the two alleles. We used SAT rather than VAT epigenomics because only SAT genome-wide epigenetic profiles are available for adipose tissue. However, analysis of the GTEx RNA-seq database (12) indicated that 87% of the 2,245 genes preferentially expressed in VAT were also preferentially expressed in SAT (ratio of ≥2.0 for VAT or SAT TPM, transcripts per million, vs. the median TPM of 35 heterologous tissues).

**Figure 1.**
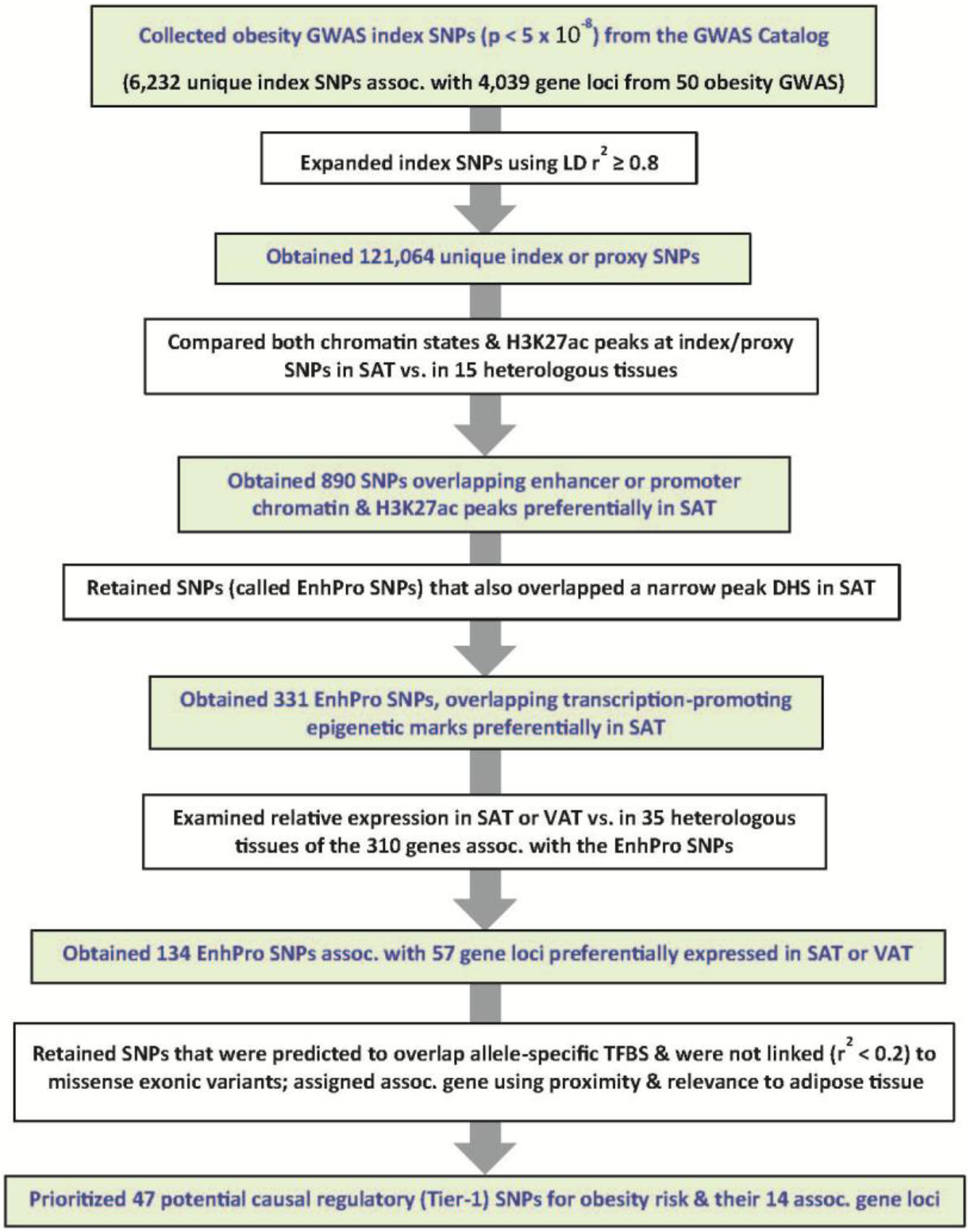
Workflow for prioritizing potential causal regulatory (Tier-1) SNPs from 50 obesity GWAS. Summary of prioritization of plausible regulatory obesity SNPs. Assoc, associated; LD, linkage disequilibrium; H3K27ac, histone H3 lysine-27 acetylation (an epigenetic mark of active enhancers or promoters); SAT, subcutaneous adipose tissue; DHS, DNaseI hypersensitive site; TFBS, transcription factor binding site(s).

From the 121,064 I/P SNPs, 331 SNPs (called EnhPro SNPs) were found to overlap a peak of open chromatin in SAT and strong enhancer or strong promoter chromatin as well as a peak of active positive regulatory chromatin (histone 3 lysine 27 acetylation, H3K27ac) preferentially in SAT (in SAT and no more than four of 15 heterologous tissues; Supplementary Table S2). Overlap of a candidate regulatory SNP with a peak of open chromatin (DNaseI hypersensitive site, DHS) in a relevant tissue increases the likelihood that a TF binds to the SNP-containing oligonucleotide sequence in that tissue. Enhancer or promoter chromatin determinations were based upon enrichment of histone H3 K4 trimethylation (H3K4me3) and H3K27ac for promoter chromatin and of H3K4me1 and H3K27ac for enhancer chromatin (RoadMap Epigenome Project (8)). We identified the subset of 134 EnhPro SNPs at 57 gene loci that are preferentially expressed in SAT or VAT by examining RNA-seq profiles from poly(A)^+^ RNA (GTEx database (12)). The definition used for preferential expression was that the ratio of SAT or VAT TPM to the median TPM of 35 heterologous tissues was >2.0 and the SAT or VAT TPM was >2.0. Forty-nine of these genes were preferentially expressed in SAT and 73% of them (36) were also preferentially expressed in VAT while eight genes were preferentially expressed in VAT but not SAT (Supplementary Table S3).

We subsequently selected those SNPs with strong predictions of overlap with allele-specific TFBS and without appreciable linkage (LD *r*^2^< 0.2, EUR) to missense SNPs. Lastly, we verified SAT-preferential expression/epigenetics of the SNP-associated gene in the UCSC Genome Browser (13). The resulting 47 SNPs are referred to as Tier-1 SNPs and were associated with 14 gene loci (Supplementary Table S4).

We retained 7 of these 14 gene loci for further study, which were linked to 18 Tier-1 SNPs, including three that were obtained by imputation (Table 1). Most of the TFs predicted to bind to these 18 Tier-1 SNPs with allele-specificity have some known adipose-related functionality (Supplementary Table S5). Seven of the 14 genes were excluded from this study because they were previously examined for GWAS regulatory SNPs (*TBX15* (14)) or because they formed a distinct functional group that will described in a separate study (extracellular matrix-related genes *COL4A2, EFEMP1, FBN1, NID2,* and *ABLIM3*). In addition, a *HOXC* gene subcluster which had nine Tier-1 SNPs in the regions of *MIR196A2, HOXC9, HOXC4, HOXC5,* and *HOXC6* was also excluded. The *HOXC4/C5/C6* intragenic Tier-1 SNPs are in moderate LD (*r*^2^ = 0.45 – 0.63) with the *MIR196A2-*overlapping SNP rs11614913 (Supplementary Table S4), a SNP that affects, in *cis*, the processing of pre-miR-196a2 (15) and was previously implicated in adipose biology (16, 17). The obesity association of the *HOXC4/C5/C6* Tier-1 SNPs might be due to their linkage with rs11614913 as determined by conditional and joint analysis (COJO (18, 19) (Supplementary Table S6).

**Table 1.** Eighteen prioritized candidates for regulatory obesity-risk variants (Tier-1 SNPs) associated with seven SAT-related genes^a^

| Gene | Tier-1 SNP for obesity association | Index proxy (I/P) or imputed (Imp) SNP | Alleles (Ref or Alt) | Obesity increasing allele (Freq EUR)<sup>b</sup> | Chromatin state at the SNP in SAT<sup>c</sup> | Gene location of SNP relative to gene | Predicted TF binding<sup>d</sup> |  | Type of additional statistical analysis that predicts assocn.<sup>e</sup> |
| --- | --- | --- | --- | --- | --- | --- | --- | --- | --- |
|  |  |  |  |  |  |  | Ref allele | Alt allele |  |
| CABLES1 | rs4445996 | Imp | A/G | G (0.91) | Str enh | 5' to TSS 1 | DNTTIP1 | MAX | None |
| " | rs9966093 | Imp | C/T | T (0.91) | Str enh | 5' to TSS 1 | ZXDA, ZXDB | RXRA | None |
| " | rs4308051<sup>*†</sup> | I/P & Imp | T/G | G (0.79) | Str enh | Near TSS 2 | BHLHE40/41 | KLF4 | None |
| " | rs28589524<sup>*†</sup> | I/P | G/A | NA (0.79) | Str enh | Near TSS 2 | NFKB1, RELA | SMAD2 | None |
| FAM13A | rs10012045<sup>*†</sup> | I/P | C/T | NA (0.56) | Str enh | Intron 2<sup>f</sup> | EPAS1 | None | None |
| " | rs2869949<sup>*†</sup> | I/P | A/G | NA (0.51) | Str enh | Intron 1 | ZNF333 | None | SMR&HEIDI |
| " | rs11097198<sup>*†</sup> | I/P | C/T | NA (0.51) | Str enh | Intron 1 | STAT1/3/4/5B | None | SMR&HEIDI |
| " | rs6532080<sup>*†</sup> | I/P | A/C | NA (0.56) | Str enh | Intron 1 | SOX4 | None | None |
| " | rs13133548<sup>*†</sup> | I/P & Imp | G/A | A (0.48) | Str enh/prom | Intron 1 | None | POU3F3 | PAINTOR; SMR&HEIDI |
| " | rs7660000 | I/P & Imp | C/T | C (0.70) | Str enh | 5' to TSS 1 | ZBTB44 | None | None |
| HOXA11 | rs530375207 | I/P | T/A | NA (0.91) | Str enh | 3'-UTR | ZNF254 | PRRX2, DLX5 | None |
| PC | rs17147932 | I/P & Imp | G/T | T (0.05) | Str enh | Near TSS 2 | KLF2/6, SP2/3 | None | None |
| PEMT | rs8078513<sup>*</sup> | I/P & Imp | C/A | A (0.07) | Str enh | End of intron 2 | SMAD3 | None | None |
| " | rs8070432<sup>*</sup> | Imp | T/C | C (0.07) | Str enh | Near TSS 3 | None | RUNX1 | None |
| RP11-392O17.1 | rs7555150<sup>*†</sup> | I/P & Imp | A/G | G (0.61) | Str enh/prom | Near prom | None | NFIB | None |
| " | rs12025363<sup>*†</sup> | I/P & Imp | G/A | A (0.61) | Str enh | Prom region | GTF2IRD1 | None | None |
| ZBED3-AS1 | rs7732130 | I/P & Imp | G/A | G (0.33) | Str enh | Near 3' end | RFX7 | None | None |
| " | rs9293708<sup>†</sup> | I/P & Imp | C/T | C (0.43) | Str enh | Near 3' end | SOX18 | None | PAINTOR |

<sup>a</sup> Tier-1 SNPs were obtained from 50 GWAS (Fig. 1); SAT, subcutaneous adipose tissue.<sup>b</sup> Obesity-increasing allele was from WHR<sub>adjBMI</sub> summary data by Pulit et al. (20); freq EUR, frequency in the European population; NA, not available (Ref allele frequency in EUR is given instead).<sup>c</sup> Str prom, strong promoter chromatin; Str enh/prom, strong promoter or enhancer chromatin; Str enh, strong enhancer chromatin from Roadmap database 18-state chromatin segmentation maps.<sup>d</sup> Transcription factors (TFs) predicted by stringent criteria to bind to SNP-containing sequences in an allele-specific manner.<sup>e</sup> The indicated Tier-1 SNP was prioritized by SMR & HEIDI or PAINTOR programs (Supplementary Tables S7 and S9); assocn, association.<sup>f</sup> Intron nomenclature for *FAM13A* is for the main isoform generated from transcription initiation at the proximal TSS.<sup>\*</sup>, the Tier-1 SNP is in LD $r^2 > 0.75$ (EUR) with other Tier-1 SNPs associated with the same gene; other multiple Tier-1 SNPs for a gene are in low LD with each other ( $r^2 < 0.3$ ).<sup>†</sup>, the Tier-1 SNP has LD $r^2 \geq 0.4$ (EUR) with 12 prioritized plausible causal SNPs ( $PP \geq 0.6$ ) from PAINTOR WHR<sub>adjBMI</sub> with epigenetic annotations (Supplementary Table S9).

### Limited power in finding best candidate regulatory SNPs with several commonly used statistical methods

As alternatives to our comparative epigenomic/transcriptomic analysis for obesity SNP prioritization (Figure 1), we tried several popular statistical methods using the largest available obesity GWAS summary data (20), some epigenetic data for SAT (8), and/or SAT eQTLs (12). First, to identify SNPs and genes with potential causal effects on obesity risk, we used the summary data-based Mendelian randomization with heterogeneity in dependent instruments test (SMR and HEIDI (9)) to prioritize SNPs that have statistically significant potential to affect both the transcript level of a gene and obesity based upon data for SAT eQTLs at a locus (12) and obesity GWAS data. The resulting 58 SNPs that were associated with 65 genes contained no SAT EnhPro SNPs, and only nine genes, including *FAM13A*, showed preferential expression in SAT (Supplementary Table S7). Only 10 of these 58 SNPs overlapped a narrow peak of DNaseI hypersensitivity in SAT and only 30 were in a DNaseI hypersensitive region in any of 53 examined tissues or cell types. Expansion of the 58 SNPs using LD *r*^2^ ≥ 0.80 (EUR) yielded 5,395 SNPs. Only three of these 5,395 SNPs (*FAM13A* SNPs rs11097198, rs13133548, and rs2869949; Table 1 and Supplementary Table S4) were Tier-1 SNPs. In contrast, 30 Tier-1 SNPs were derived by our epigenomic/transcriptomic scheme from Pulit *et al.* GWAS summary data (20).

We also did colocalization analysis (eCAVIAR (21)), to look for potential causal variants at a given locus that could be causal in both GWAS and eQTL studies among the seven studied gene regions (Table 1). None of our EnhPro/Tier-1 SNPs nor any new credible regulatory SNPs were prioritized by colocalization analysis (Supplementary Table S8). In addition, we conducted fine-mapping analysis (Probabilistic Annotation INtegraTOR, PAINTOR (22)), to prioritize potential causal SNPs from the neighborhoods of the above-mentioned seven genes. The fine-mapping analysis of WHR_adjBMI_ that included SAT epigenetic parameters prioritized only two of 18 Tier-1 SNPs (*FAM13A* rs13133548, posterior probability, PP =1, and *ZBED3-AS1* rs9293708, PP = 0.8; Supplementary Table S9). Of the other 55 SNPs prioritized by this fine-mapping analysis (PP ≥ 0.6), only four overlapped enhancer or promoter chromatin and a peak of DNaseI hypersensitivity in SAT (*FAM13A*: rs4544678, rs3775380; *HOXA11*: rs17471520: and *PEMT*: rs8070128). Importantly, four of the seven gene loci had at least one fine-mapping prioritized SNP which was in moderate to high LD (*r*^2^ > 0.4) with a Tier-1 SNP (Table 1 and Supplementary Table S9). Although regulatory SNPs can modulate obesity through tissues other than SAT, Tier-1 SNP rs7732130, which as described below was implicated in T2D risk in pancreas by detailed experimental analysis (23, 24), was not prioritized by any of the analytical methods that we used, even those that did not include SAT epigenetic annotation (Supplementary Table S7-9).

Because the protocol used in the present study to obtain Tier-1 SNPs (Figure 1) was not meant to be all-inclusive, but rather to give some highly credible risk alleles, we tried to expand it. The last criterion for being a Tier-1 SNP, overlap of a stringently predicted allele-specific TFBS, is limited in power. Five EnhPro SNPs associated with two of the seven genes in Table 1, *FAM13A* and *RP11-392O17.1,* did not meet this last criterion. COJO analysis (18, 19) to look for secondary association signals for the Tier-1 SNPs at neighboring *FAM13A* and *RP11-392O17.1* EnhPro SNPs revealed three *FAM13A*-associated EnhPro SNPs (i.e., rs17014602, rs4544678 and rs3775378) that might still be credible obesity risk-associated variants. The p-value for the *FAM13A* Tier-1 SNP rs13133548 increased after conditioning on these three EnhPro SNPs (Supplementary Table S6). Therefore, our protocol could be extended in the future to include COJO analysis of EnhPro SNPs that did not have predictions of overlapping allele-specific TFBS.

### *CABLES1, PEMT,* and *PC* genes are associated with obesity Tier-1 SNPs that may regulate alternate usage of transcription start sites

One or more of the Tier-1 SNPs associated with the genes encoding Cdk5 And ABL1 Enzyme Substrate 1 (*CABLES1*), Phosphatidylethoanolamine N-Methyltransferase (*PEMT*), and Pyruvate Carboxylase (*PC*) were located within 0.7 kb of an alternative transcription start site (TSS; Figure 2A and Supplementary Figures S1-S5). *CABLES1, PEMT,* and *PC* are the only genes in their 1-Mb neighborhoods that exhibit preferential expression in SAT (Figure 2 and Supplementary Table S10). However, expression of these genes is not specific for adipose tissue, and correspondingly, they have non-adipose tissue-specific functions and generalized functions as well as adipose-related functions (25). Although *CABLES1* is involved in the regulation of proliferation of pituitary gland cells (26), its expression in SAT or VAT is about 3 times that in the pituitary gland (Supplementary Table S10).

**Figure 2.**
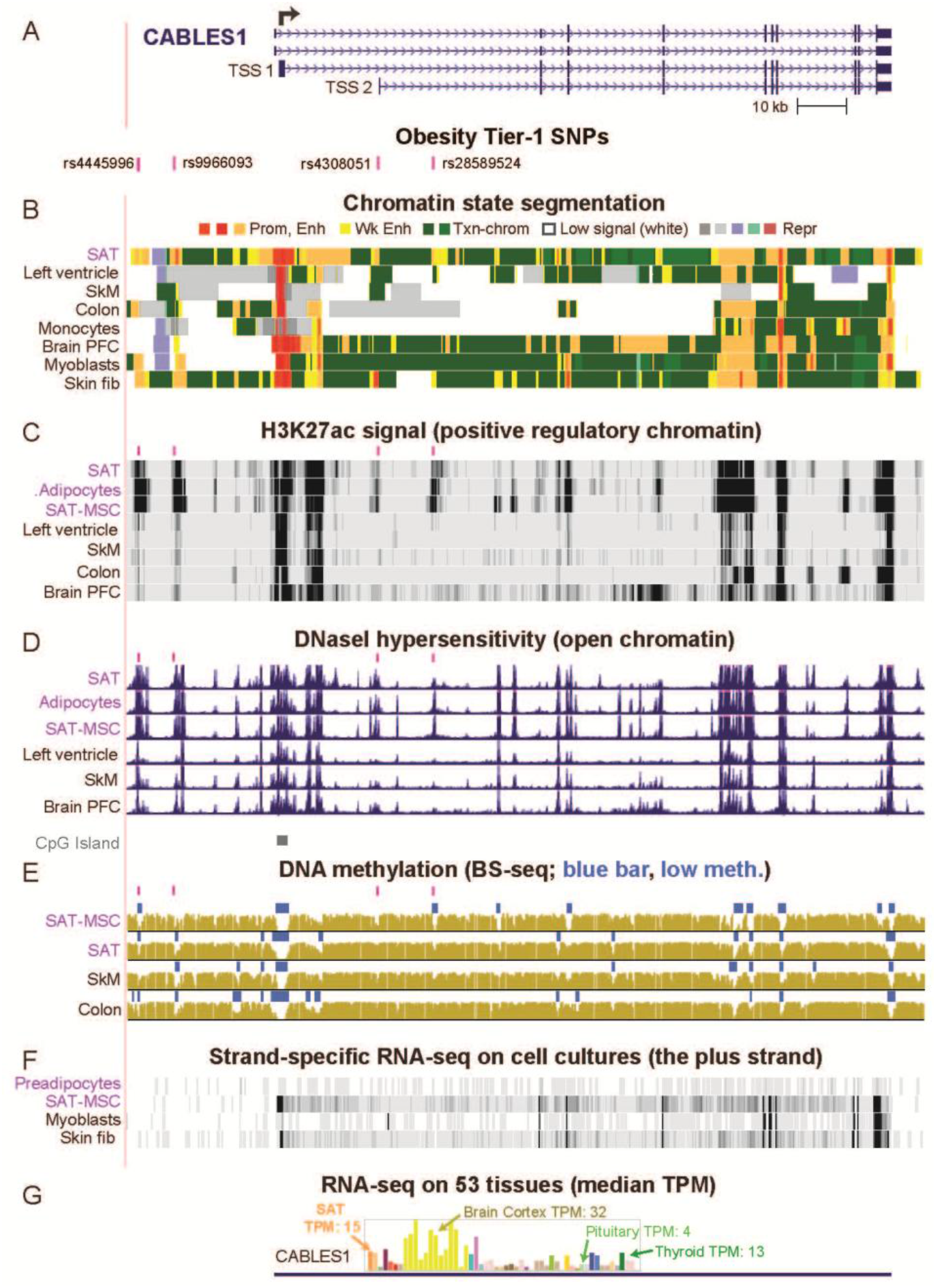
*CABLES1*, which controls cell growth, is associated with Tier-1 SNPs that might regulate alternative usage of its widely separated TSS. **A**. A 162-kb region containing *CABLES1* and four prioritized regulatory SNP candidates (Tier-1 SNP) for obesity risk (chr18:20,684,550-20,847,013). The broken arrow denotes the most frequently used TSS. **B**. Strong promoter (Prom) or strong enhancer (Enh) chromatin segments; weak enhancer chromatin (Wk Enh); chromatin with the H3K36me3 mark of actively transcribed regions (Txn-chrom); chromatin with low signals for the assessed histone modifications; repressed (Repr) chromatin (8). **C**. H3K27ac enrichment profiles (8). **D**. Profiles of open chromatin (DNaseI hypersensitivity (8)) and a track for CpG islands, CpG-rich regions (13). **E**. DNA methylation profiles from whole-genome bisulfite sequencing; blue horizontal bars indicate regions of significantly low DNA methylation relative to the rest of the same genome. **F**. RNA-seq on cell cultures (13); just the plus strand is shown with a vertical viewing range of 0 – 100. **G**. Tissue RNA-seq depicted as a bar graph with median TPM from hundreds of biological replicates (12). First two orange bars, SAT and VAT; yellow bars, brain samples; see Supplementary Table S10 for details. SAT, subcutaneous adipose tissue; SkM, skeletal muscle; PFC, pre-frontal cortex; fib, fibroblasts; MSC, mesenchymal stem/stromal cells; short pink bars in panels **C**, **D**, and **E**, indicate positions of Tier-1 SNPs. All tracks were visualized in the UCSC Genome Browser (hg19) and are aligned in this figure and Figures 3–5.

There were tissue-specific differences in the usage of alternative TSS for *CABLES1, PEMT,* and *PC*, including in SAT vs. some other tissues (Supplementary Figures S2, S4, and S5), that might be influenced by Tier-1 SNP-containing enhancer chromatin near one of the TSS. Enhancer chromatin, peaks of H3K27ac, and DNaseI hypersensitivity overlapped some of the Tier-1 SNPs for these genes not only preferentially in SAT but also in SAT-derived mesenchymal stem/stromal cells (SAT-MSC) or adipocytes derived from them *in vitro* (27) (Table 1, Figure 2B - D; Supplementary Figures S1, S3, and S5). DNA hypomethylated regions, a frequent hallmark of *cis*-acting positive transcription-regulatory elements, overlapped several of *CABLES1* Tier-1 SNPs specifically in SAT-MSC, adipocytes, brain neurons, and the nonneuronal cell fraction of brain (Figure 2E and Supplementary Figure S1). The brain cell-specific hypomethylation is likely to reflect specific roles for *CABLES1* in the brain (28) and might be related to differences in TSS usage in brain vs. adipose (Supplementary Figure S2). The Tier-1 SNP-containing enhancer chromatin of *CABLES1, PEMT,* and *PC* may regulate overall transcription as well as TSS usage. This is illustrated by melanocytes, which had very high overall levels of expression of *CABLES1*, extensive enhancer chromatin and H3K27ac around TSS 2, and much transcription initiation from TSS 2 as well as TSS 1 (Supplementary Figures S1 and S2B and data not shown).

The tissue-specific TSS usage for *CABLES1* and *PEMT* results in very different sized polypeptides (Supplementary Figures S2B and S4C). However, despite the tissue-specificity for alternate TSS usage for *PC* (*ENSG00000173599*; Supplementary Figure S5F), there is no effect of TSS choice on the encoded polypeptide because the open reading frame (ORF) starts far downstream of all TSS. In addition to the annotated TSS 2, we found a novel, cell type-specific TSS (TSS 3) that is even closer to Tier-1 SNP rs17147932 (0.7 vs. 2.5 kb upstream; Supplementary Figure S6). Only transcription initiation from TSS 1 allows a *PC*-intragenic *MIR3163* gene to be included in the primary transcript (Supplementary Figure S5A). The mature miR-3163 has many documented effects on cell biology partly through modulation of Wnt signaling (29). SAT and liver share promoter/enhancer chromatin, a H3K27ac peak, and a DHS overlapping rs17147932 at *PC* (Supplementary Figure S6). The frequency of TSS usage might be modulated by this Tier-1 SNP in liver, which has the highest expression of *PC,* and in adipose, which has the second highest expression level. Supporting a role of TSS 2 in both the adipose and liver lineages, a binding site in adipocytes was found at the core promoter region of TSS 2 for the adipogenesis-associated TF PPARG (30) and in liver and HepG2 cells at the core promoter regions of TSS 2 and TSS 3 for many transcription initiation-associated proteins (TF ChIP-seq; Supplementary Figure S6).

### Only one of the previously prioritized *FAM13A* candidate regulatory SNPs for obesity risk is among the six Tier-1 SNPs identified for this gene in the present study

*FAM13A* (*Family With Sequence Similarity 13 Member A*) was associated with six Tier-1 SNPs (Figure 3A, Table 1), four of which were found to be eQTLs in SAT (Supplementary Table S11). This gene is implicated in modulating diet-induced obesity, regulating insulin signaling in adipocytes, participating in Wnt signaling, and affecting the risk of certain lung diseases (3, 31, 32). We evaluated five previously reported candidate regulatory SNPs (Figure 3A, blue bars and blue font). One, rs2276936, was highlighted by Lin *et al*. (33) as a likely regulatory SNP from massively parallel reporter assays (MPRA) on bronchial epithelial cells, GWAS data for body fat distribution and blood lipid levels, reporter gene experiments on HepG2 cells, and a CRISPR/Cas-9 mediated ~0.1-kb deletion around this SNP in HepG2 cells. We found that the 0.5-kb region centered around rs2276936 (Figure 3A, blue arrow) did not overlap a peak of DHS (open chromatin) or of H3K27ac in >15 types of cells or tissues, including SAT, liver, HepG2 cells, lung, and esophagus, although the SNP was embedded in a broad region of enhancer chromatin in SAT (Figure 3A-D). Therefore, rs2276936 does not meet the general expectation that a SNP whose allelic state modulates transcription is located at a TF-binding site within (and not just near) a peak of open chromatin and of H3K27ac in a biologically relevant cell type or tissue.

**Figure 3.**
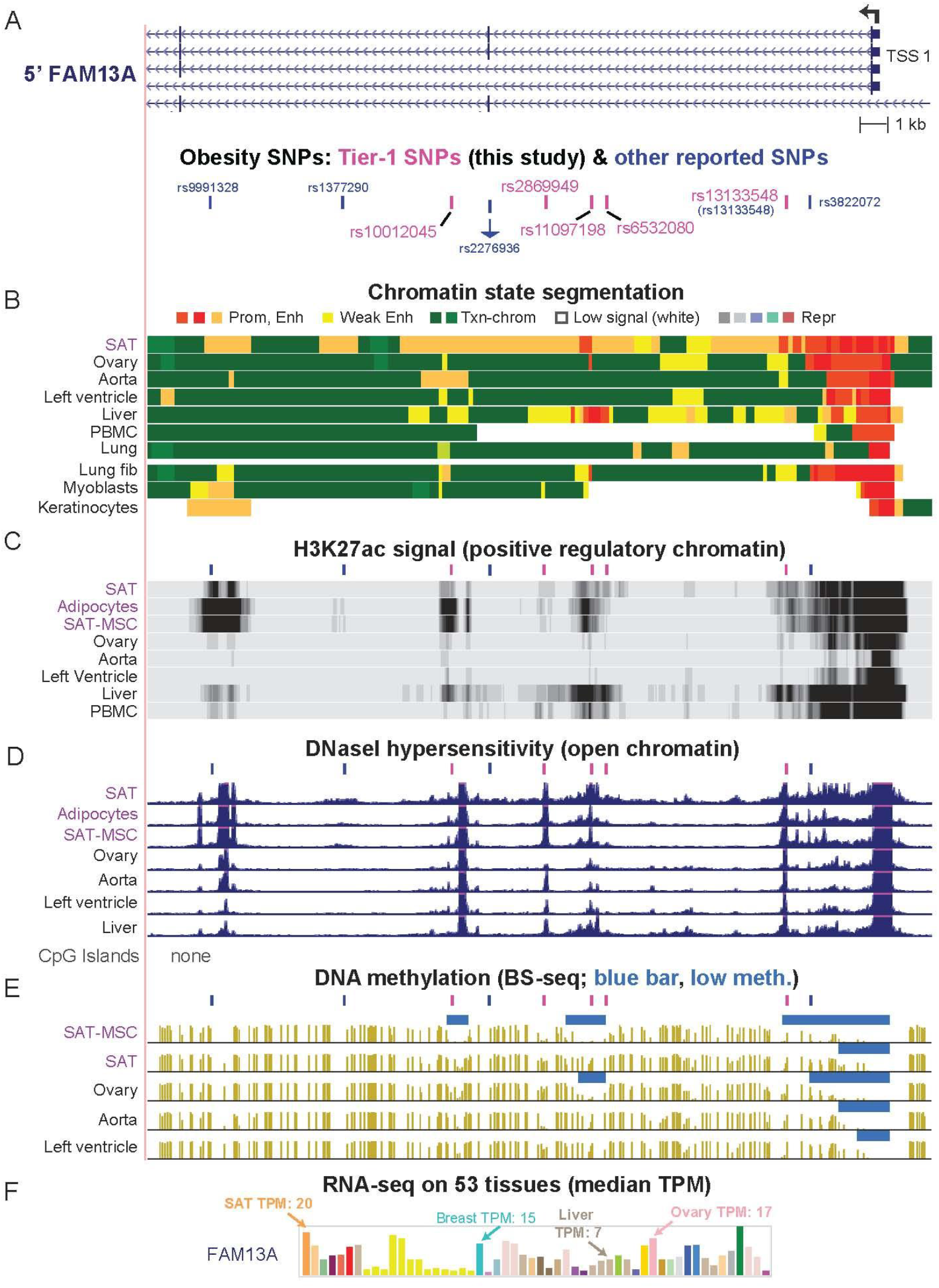
One of the previously highlighted obesity-related SNPs in *FAM13A* (rs13133548) has optimal epigenetic features to be a obesity-risk regulatory SNP. **A**. A 37-kb region (chr4:89,710,207-89,746,974) overlapping five of the six Tier-1 SNPs (pink bars) as well as four other SNPs that had been described as regulatory SNP candidates from obesity GWAS (blue bars); rs13133548 is the one Tier-1 SNP that was previously described. **B**, **C**, **D,** and **E**. Chromatin state segmentation, H3K27ac, DHS, and bisulfite-seq profiles as in Figure 2. **F**. Bar graph of median expression levels for different tissues as in Figure 2. Pink or blue bars above tracks in panels **C**, **D**, and **E**, indicate positions of Tier-1 SNPs and previously highlighted candidates for regulatory SNP, respectively. PBMC, peripheral blood mononuclear cells.

Of the other four other previous regulatory candidate SNPs in this region (3, 11, 34), rs1377290 did not overlap a DHS peak or a H3K27ac peak in SAT and rs9991328 did not overlap a DHS peak in SAT or adipocytes (Figure 3A-D). Another of these previously reported SNPs, rs3822072, lacked specificity for its peak of H3K27ac (present in eight of 15 non-adipose tissues) and was missing a predicted overlapping allele-specific TFBS. The fourth SNP, rs13133548 (35), met all the criteria for a Tier-1 SNP. These five previously described regulatory candidate SNPs are in high or perfect LD (*r*^2^ = 0.78 – 1.00, EUR) with our four novel Tier-1 SNPs in introns 1 or 2 of the short isoforms of *FAM13A* (Table 1). The last novel Tier-1 SNP, rs7660000, is ~7 kb upstream of TSS 1 in intron 7 of the long *FAM13A* isoform initiated at TSS 2 and located 235 kb upstream of TSS 1 (Supplementary Figure S7). Only the long FAM13A protein isoform (Supplementary Figure S7) contains a signaling-associated RhoGAP domain (36). More transcript is made from *FAM13A* TSS 1 than TSS 2 in SAT, SAT-MSC, and preadipocytes as well as most, but not all cell types (Supplementary Figure S7). Tier-1 SNPs for *FAM13A* might contribute to different frequencies of use of TSS 1 and TSS 2 as well as tissue-specific differences in expression levels of this gene (Figure 3F).

### *RP11-392O17.1*, an often-overlooked SAT-specific lncRNA gene, has novel Tier-1 SNPs in its promoter region

Of the 14 gene loci associated with Tier-1 SNPs, only *RP11-392O17.1, HOXA11, MIR196A2/HOXC4*,*5,6,9* and *TBX15* displayed a strong bias for expression (12) in SAT vs. VAT as well as for SAT vs. non-adipose tissues (Figure 4A, Supplementary Table S3). As described above, we did not further examine the *HOXC* and *TBX15* loci in this study. For *HOXA11,* a single Tier-1 SNP in the 3’ untranslated region (3’-UTR) near both *HOXA11-AS1* and *HOXA10* was identified (Supplementary Figure S8). The 3’ location of this SNP raises the possibility that it is involved in post-transcriptional control. In contrast, rs7555150 and rs12025363, the novel Tier-1 SNPs for *RP11-392O17.1,* are 0.6 or 2.3 kb upstream of the TSS in promoter or enhancer chromatin in SAT, which suggests a possible transcription regulatory function. These SNPs are in perfect LD with each other and overlap SAT eQTLs for *RP11-392O17.1* (Supplementary Table S11), as reported previously for other nearby SNPs (37). The *RP11-392O17.1* Tier-1 SNPs display a much stronger association with female rather than male obesity (38), and gender bias was also found for *FAM13A, PEMT,* and *ZBED3-AS1* Tier-1 SNPs (Supplementary Tables S4).

**Figure 4.**
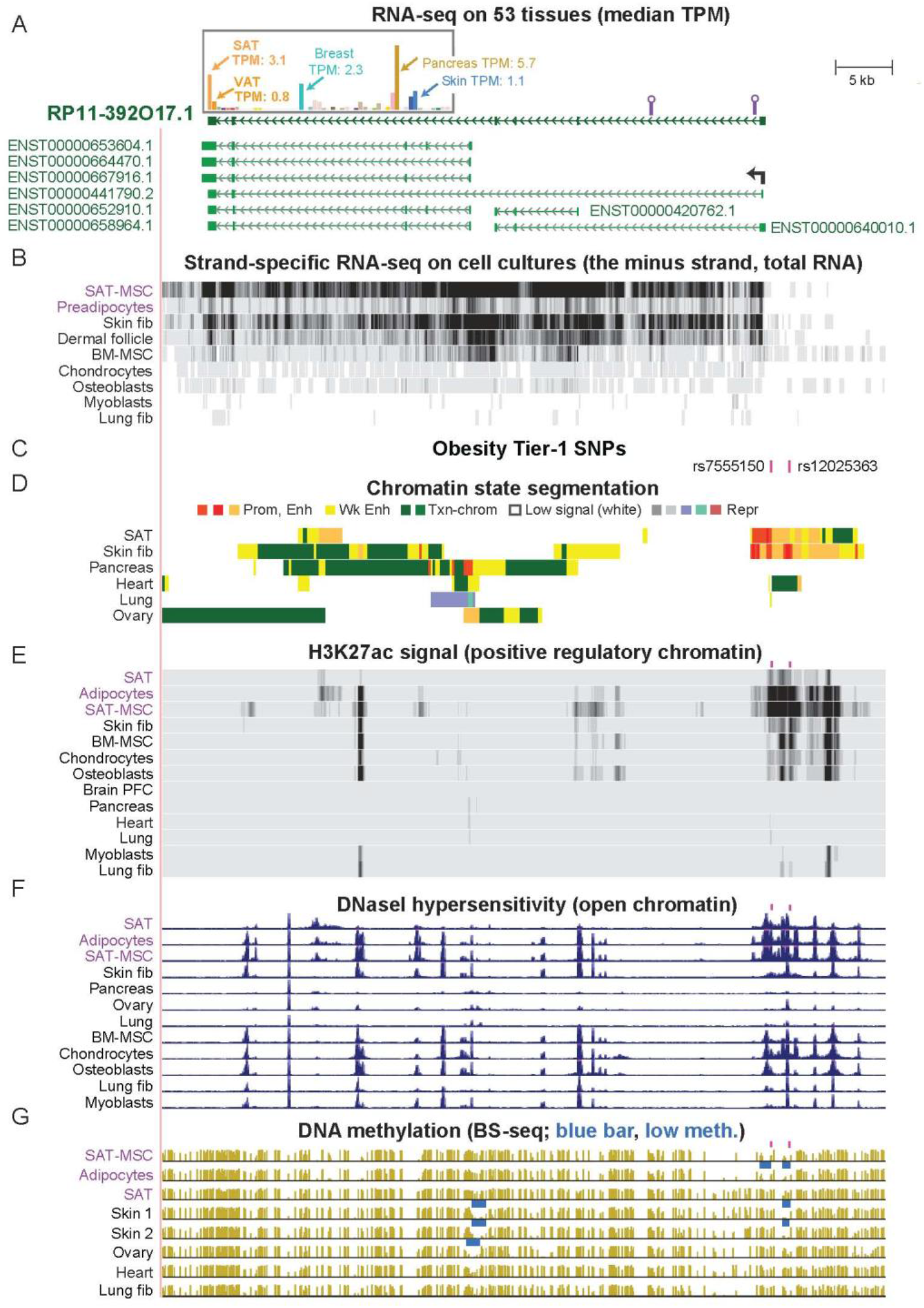
A lncRNA gene, *RP11-392O17.1*, associated with SAT-related and female obesity that is often overlooked in assignments of genes to obesity GWAS SNPs. **A**. A 65-kb region (chr1:219,578,827-219,643,464) containing *RP11-392O17.1*; above it is the tissue RNA-seq profile (GTEx, determined from poly(A)^+^ RNA) for the gene structure depicted immediately underneath it. The *RP11-392O17.1* isoforms shown are from the latest Ensembl Comprehensive Gene Annotation Set (v104) but this gene is not yet in the RefSeq database. Purple lollipops, regions that we amplified by RT-PCR as described in the text. **B**. Cell culture RNA-seq (showing the minus strand) on total RNA, vertical viewing range 0 – 50. **C**. The location of two Tier-1 SNPs. **D**, **E**, **F**, and **G.** Chromatin state segmentation, H3K27ac, DHS, and bisulfite-seq, as in the previous figures. There were no CpG islands in the region shown. BM-MSC, bone marrow-derived MSC.

*RP11-392O17.1* is a little-studied lncRNA gene, which has recently been implicated in adipogenesis (39). The 3-Mb region surrounding this gene has no coding gene with preferential transcription in SAT vs. VAT, as does *RP11-392O17.1* (Supplementary Table S10). However, the very small and very weakly expressed lncRNA gene *RP11-95P13.2* (SAT TPM, 0.5) 98 kb upstream of *RP11-392O17.1* does show preferential expression in SAT. *RP11-392O17.1* has not yet been designated a RefSeq gene although it is in the Ensembl gene database (*ENSG00000228536*) and displays highly cell type-specific expression in SAT-MSC, SAT, and adult skin fibroblasts (Figure 4B and Supplementary Figures S9 and S10A). This gene had transcription-promoting epigenetic marks specifically in SAT, adipocytes, and SAT-MSC in the region of its Tier-1 SNPs, rs7555150 and rs12025363 (Figure 4C-G). Moreover, 13 bp downstream of its TSS, there is a binding site in adipose stem cells for, the adipose-associated PPARG TF (Unibind database (13, 40)).

In 13 GWAS related to obesity ((41, 42) and Supplementary Figure S9 legend), many candidate regulatory SNPs located 11 – 118 kb upstream of *RP11-392O17.1* were previously associated with *LYPLAL1* (*Lysophospholipase Like 1*) or *LYPLAL1-AS1. LYPLAL1* is broadly expressed and located 0.2 Mb further from these SNPs than is its closest downstream gene, *RP11-392O17.1*. The RefSeq gene structure *LYPLAL1-AS1* (NR_135822.1), which partly overlaps *RP11-392O17.1,* displayed no evidence for transcription in any of >20 examined cell types (Supplementary Figures S9 and S10) and is not included in the GTEx database for tissue expression. In addition, chromatin state segmentation tracks show the absence of promoter chromatin at the 5’ end of the RefSeq *LYPLAL1-AS1* in tissues and cell cultures unlike its presence at the 5’ end of *RP11-392O17.1* in SAT and adult skin fibroblasts (Figure 4D and Supplementary Figure S10B). Nonetheless, the name *LYPLAL1-AS1* is sometimes used in the literature to denote *RP11-392O17.1* or *LYPLAL1-DT,* a lncRNA gene upstream of *LYPLAL1* (Supplementary Figure S9). For example, Yang *et al.* (39) referred to a 0.5-kb region from *RP11-392O17.1* (Supplementary Figure S9A, blue box) as *LYPLAL1-AS1.* Importantly, these previously identified obesity GWAS SNPs exhibited moderate LD (*r*^2^ = 0.35 to 0.68, EUR) with Tier-1 SNPs in the promoter region of *RP11-392O17.1*. Therefore, they may have been detected in obesity GWAS only because of their LD with *RP11-392O17.1* promoter-region SNPs for this gene.

We tried to test for allele-specific expression of *RP11-392O17.1* in a SAT-MSC cell strain that we identified as heterozygous for both Tier-1 SNPs. Using RT-PCR on total RNA, we amplified two small regions of cDNA not far from the 5’ end of the gene (Figure 4A, lollipops) that contained SNPs in high LD with our Tier-1 SNPs. Sanger DNA sequencing of the PCR products (Supplementary Figure S11) from the cDNA and from the analogous genomic DNA revealed amplified background sequences that interfered with quantification of the relative expression of the alleles. We were only able to verify transcription of the examined 5’ portions of the gene in SAT-MSC from both alleles, unlike the finding on adipose tissue from reanalysis of RNA-seq data (43).

### Two obesity Tier-1 SNPs located in *ZBED3-AS1* might help control expression in SAT of several genes in *cis*

We identified Tier-1 SNPs rs7732130 and rs9293708 in the 3’ end of the lncRNA gene *ZBED3-AS1* (Figure 5A). *ZBED3-AS1* might influence expression in SAT of several genes in its 1-Mb neighborhood, including *ZBED3*, which encodes Zinc Finger BED Domain-Containing Protein 3 and is oriented head-to-head with *ZBED3-AS1* with a 0.5 kb overlap*. ZBED3* and *ZBED3-AS1* have been implicated in adipogenesis (44). The *ZBED3-AS1* Tier-1 SNPs are in low LD with each other (*r*^2^ = 0.2, EUR) and reside 5 kb apart in separate enhancer chromatin segments and DNase hypersensitivity peaks in SAT (Figure 5B-D). One of them, rs9293708, is located 0.6 kb from a binding site for CTCF (CCCTC-binding factor), which can enable long-distance chromatin looping (Figure 5E and F). Despite *ZBED3-AS1* and *ZBED3* sharing a bidirectional promoter, there is appreciable *ZBED3-AS1* RNA in only one of 19 examined cell strains (melanocytes) in contrast to the broad expression of *ZBED3* in cell cultures (Figure 5G and Supplementary Figure S12). Among tissues, *ZBED3-AS1* and *ZBED3* were preferentially expressed in thyroid, SAT and VAT, as were *ZBED3-AS1* neighbors *PDE8B* (encoding Phosphodiesterase 8B) and *CRHBP* (encoding Corticotropin Releasing Hormone-Binding Protein; Figure 5H and Supplementary Table S10 for *CRHBP*). The latter two genes are relevant to obesity. *PDE8B* is involved in insulin signaling in the pancreas and is implicated in controlling lipolysis and the ratio of VAT/total adipose tissue (45, 46). Overexpression of *CRHBP* can affect weight gain in mice (47). *ZBED3-AS1* was selectively transcribed *in vivo* in adipocytes as well as in SAT, as seen in a single-nucleus RNA-seq analysis of tissues (48) (Supplementary Figure S13).

**Figure 5.**
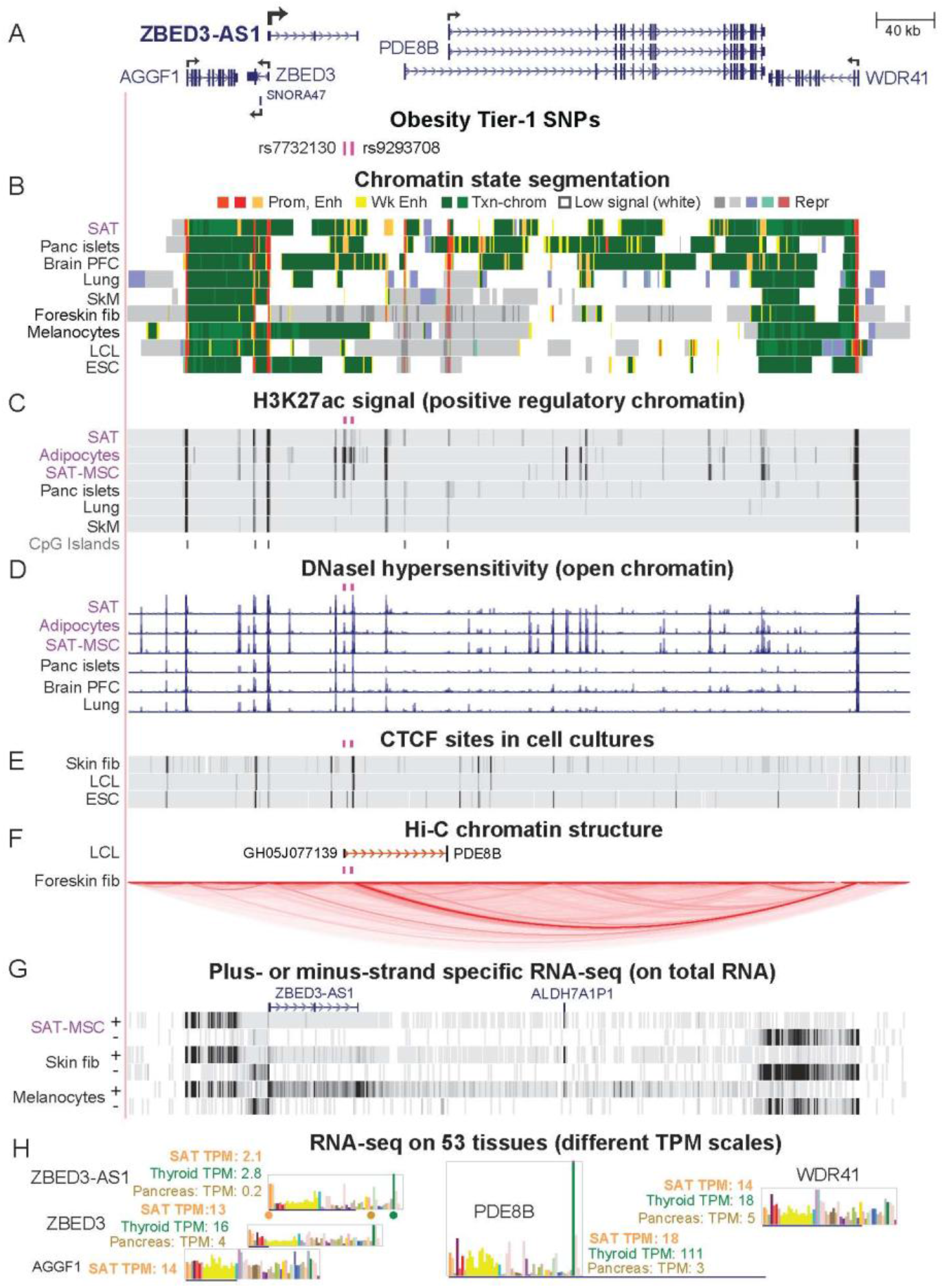
Two Tier-1 SNPs in a lncRNA gene, *ZBED3-AS1*, and neighbors *ZBED3, SNORA47, PDE8B*, and *WDR41* are near long-distance chromatin looping subregions. **A.** A 538-kb region (chr5:76,285,728-76,823,921) containing *ZBED3-AS1* and neighboring genes; for clarity, additional isoforms of *ZBED3* and *PDE8B* are shown in Supplementary Figure S14. The main TSS for *PDE8B* is indicated by a broken arrow. *CRHBP*, which is referred to in the text but not shown, is 70 kb upstream of *AGGF1*. **B, C**, and **D**. Chromatin state segmentation, H3K27ac, and DHS as in the previous figures. **E**. CTCF binding by ChiP-seq profiling, for several cell cultures. **F**. A promoter-capture Hi-C interaction for the lymphoblastoid cell line (LCL) GM12878 and all chromatin interactions in the region detected by Micro-C for foreskin fibroblasts. **G**. RNA-seq (showing results for transcription from both strands separately) on cell cultures with vertical viewing range 0 – 30; preadipocytes exhibited no detectable expression of *ZBED3-AS1* (not shown); *ALDH7A1P1* is a pseudogene. **H**. Tissue RNA-seq shown as a bar graph with dots color-coded to indicate their position in the bar graph for *ZBED3-AS1*. ESC, H1-embryonic stem cells. CpG islands (not shown) overlap all the TSS indicated by broken arrows.

In pancreatic islets, Miguel-Escalada *et al*. (24) found that a 0.9-kb enhancer chromatin region containing Tier-1 SNP rs7732130 interacts with and upregulates transcription from promoters of *ZBED3-AS1* and the neighboring *PDE8B, ZBED3*, *ZBED3-AS1, SNORA47* and *WDR41* genes. The ratio of expression of *ZBED3-AS1* in SAT or VAT vs. the median of 35 dissimilar tissues is 4; in contrast, the analogous ratio for pancreas is only 0.5 (Supplementary Table S10); however, expression levels may be higher in pancreatic islets for which comparable RNA-seq data are not available. Another distinction between SAT and pancreatic islets is that pancreatic islets lacked the enhancer chromatin in which the second Tier-1 SNP rs9293708 is embedded although both tissues had enhancer chromatin overlaying rs7732130 (Figure 5A-C; Supplementary Figure S12).

Chromatin interaction data for this region were not available for adipocytes. However, mapping of long-range chromatin interactions for a lymphoblast cell line (LCL, GM12878) and foreskin fibroblasts (49, 50) suggests that regions near both *ZBED3-AS1* SNPs are involved in such interactions and may be at the boundaries of topologically associating domains (TADs). In GM12878, there was evidence for an interaction between an ~60-bp subregion 0.1 kb from rs7732130, which exhibited weak enhancer chromatin, and a repressed promoter of *PDE8B* ~70 kb downstream (promoter capture Hi-C profile, Figure 5F). Foreskin fibroblasts displayed a strong and cell type-specific TAD that extended from a constitutive CTCF-binding site 0.5 kb away from rs9293708 in repressed chromatin to another constitutive CTCF binding site in the active *WDR41* promoter 342 kb away (Figure 5B, E, and F and Supplementary Figure S14). The TAD was seen in Micro-C chromatin interaction profiles, a modified type of Hi-C (51) that is not restricted to interactions using promoter regions as bait. The role of this TAD in foreskin fibroblasts is not clear but it might be needed to insulate the several *PDE8B* promoters from enhancer chromatin seen in skin fibroblasts more than 0.3 Mb upstream (data not shown). Although some of the effects on obesity risk of genes in this neighborhood are probably attributable to pancreas and the thyroid gland (for which epigenome profiles were not available), epigenomics and transcriptomics suggests that the two *ZBED3-AS1* Tier-1 SNPs act through adipose tissue to moderate inherited obesity risk.

## Discussion

We prioritized 47 candidates for obesity-risk regulatory SNPs (Tier-1 SNPs) using a detailed approach involving comparisons of epigenomics and transcriptomics in adipose tissue vs. non-adipose tissue. Of the 18 SNPs that were studied in detail, all but two (rs13133548 and rs7732130 (24, 35)) had not been previously reported as possible regulatory SNPs. We showed that identifying credible candidates for obesity-risk regulatory SNPs using their SAT epigenetics (because VAT epigenomic databases were not available) and the expression in SAT and VAT of their associated genes usually prioritized genes preferentially expressed in VAT as well as in SAT. Thus, these candidates for regulatory SNPs are likely to be relevant to obesity-related disease susceptibility (3).

Specific roles in adipose biology have been inferred for the seven gene loci associated with these SNPs (3, 5, 25, 39, 44, 52–58). Remarkably, four of these genes, *CABLES1, PEMT, PC,* and *FAM13A*, had one or more of their Tier-1 SNPs near an alternate tissue-specific TSS. The choice of alternative well-separated TSS for *in vivo* transcription of these genes is consequential because it affects either the encoded protein structure (*CABLES1, PEMT,* and *FAM13A*) or the expression of a miRNA gene in an isoform-specific intron (*PC*; Supplementary Figures S2, S4, and S5). Therefore, Tier-1 SNPs for these genes could be involved in modulating the structure of the resulting transcript and not just its quantity in SAT and, thereby, affect inherited obesity risk.

The variants at Tier-1 SNPs at *CABLES1, FAM13A, PC*, *PEMT, ZBED3-AS1, RP11-392O17.1* are highly plausible candidates for modulating transcription. The genes associated with these SNPs are involved in cell signaling, metabolism, and/or post-transcriptional regulation in ways that could modulate adipose tissue formation or physiology. *CABLES1* encodes a growth suppressor protein that acts as a signaling hub for cell growth (59). It plays a central role in regulating the cell cycle, cell proliferation, and cancer (60). Although it has not been previously directly linked to obesity, some of its regulatory interactions are very likely to impact obesity (53, 61). For example, CABLES1 affects CDK5 phosphorylation, which, in turn, controls phosphorylation of the critical adipose-related TF PPARG (52). FAM13A protein is also implicated in diverse types of signaling (62), including adipocyte insulin signaling (32) and Wnt/beta-catenin signaling, which is crucial for normal development and homeostasis of adipose tissue and other lineages (63, 64). Knockdown of *FAM13A* during *in vitro* adipogenesis increases expression of several adipocyte markers (11, 34) while *in vitro* overexpression leads to preadipocyte apoptosis (65). Although, mouse double-knockout or overexpression of this gene results in little or no change in adiposity, animals fed high-fat diets exhibited significant decreases in the VAT/SAT ratio in male *FAM13A* knockout mice (11, 31, 65).

Tier-1 SNP-associated lncRNA genes *RP11-392O17.1* and *ZBED3-AS1* have recently been implicated in adipose biology through interaction with specific proteins or RNAs. In SAT-MSC cells differentiating *in vitro* to adipocytes, overexpression of part of *RP11-392O17.1* (called *LYPLAL1-AS1* despite the lack of evidence for expression of an overlapping *LYPLAL1-AS1* gene structure; Supplementary Figure S10) increased expression of adipogenesis markers and fat droplet deposition (39). This finding was partially attributed to the stabilization of desmoplatin, a desmosome protein, and inhibition of Wnt signaling. The prominence of the active chromatin profiles of *RP11-392O17.1* in SAT-MSC and adipocytes is consistent with expression of this gene playing a role in adipocyte progenitor differentiation as well as in adipocyte function (Figure 4). Nonetheless, in 13 obesity-related GWAS (Supplementary Figure S9), many candidate risk SNPs upstream of *RP11-392O17.1* were assigned to *LYPLAL1* rather than to the much closer *RP11-392O17.1.* However, *LYPLAL1* displays no preferential expression in any obesity-relevant tissue, unlike *RP11-392O17.1*. On the basis of eQTLs and limited transcription data, in two previous studies, it was proposed that a few obesity GWAS-derived SNPs that were ascribed to *LYPLAL1* may actually be associated with *RP11-391O17.*1 (37, 38). Here, extensive transcription, epigenetic, and genetic data support such reassignment from *LYPLAL1* to *RP11-392O17.1* 0.2 Mb downstream from it, and the conclusion that the previously described SNPs in the *RP11-392O17.1*-upstream intergenic region are proxies for regulatory SNPs in the promoter region of this lncRNA gene.

*ZBED3-AS1,* which contains two intragenic Tier-1 SNPs, rs7732130 and rs9293708 (Figure 5), encodes an RNA that acted as a competitive endogenous RNA (ceRNA, sponge RNA) for miR-513a-5p and miR-381-3p (66). These miRNAs are implicated in nonalcoholic fatty liver disease (67) or gene dysregulation in VAT in morbidly obese individuals (68). *ZBED3-AS1* displays increases in its steady-state RNA levels during *in vitro* adipogenesis and levels of this RNA correlate with expression of adipogenesis markers in liposuction samples (44). Moreover, production of *ZBED3-AS1* RNA was reported to be necessary for normal levels of expression of markers of adipogenesis during *in vitro* differentiation (44). *ZBED3-AS1* poses an interesting example of a gene with intragenic enhancers that may act in *cis* on nearby genes as well as on itself. The results of Xu *et al*. (44) suggest that *ZBED3-AS1* also functions in *trans* in SAT-MSC.

In a study of T2D GWAS SNPs, Miguel-Escalada (24) documented long-distance chromatin interactions in pancreatic islets between enhancer chromatin containing rs7732130 and promoters of neighboring genes. Epigenome editing showed that these interactions can potentiate transcription of *ZBED3-AS1* gene neighbors as well as *ZBED3-AS1*. Moreover, in reporter gene assays on pancreatic islet cells, a 0.9-kb region harboring this SNP gave allele-specific upregulation (23). We independently determined that rs7732130 and nearby rs9293708 are obesity GWAS-derived Tier-1 SNPs. Our analysis combined with these two previous studies suggests that both SNPs modulate obesity risk through adipose tissue and pancreas while rs9293708, located in a separate enhancer chromatin region specific to SAT, does so only through adipose tissue. Given the higher expression of *ZBED3-AS1* in SAT than in pancreas (Supplementary Table S10), the influence of rs7732130 on obesity risk might derive more from its effect on expression of this gene in adipose (both SAT and VAT) than in pancreas.

The Tier-1 SNPs of *PC* and *PEMT* might also modulate inherited obesity risk through other tissues as well as through adipose. Positive transcription regulatory marks at the *PC* and *PEMT* intragenic Tier-1 SNPs were seen in both liver and SAT, the tissues that most highly express these genes (Supplementary Figures S3 and S5). These findings suggest that regulatory SNP candidates for obesity risk might assert their effect in more than one tissue. Previous studies indicated that many obesity GWAS regulatory SNPs even derive their obesity associations from brain, a tissue very dissimilar to adipose tissue (10). The role in brain of such regulatory SNP candidates could be evaluated by our scheme focusing on the epigenomics and transcriptomics of brain instead of adipose.

We demonstrated that a detailed epigenomic/transcriptomic/genetic approach is not only valuable for discovery of novel credible candidates for regulatory SNPs (Tier-1 SNPs) but also for re-evaluating previously reported candidate regulatory SNPs. We also showed that most of our prioritized Tier-1 SNPs were not detected with several standard statistical tools for evaluating GWAS SNPs (SMR and HEIDI (9), PAINTOR (22), and eCAVIAR (21)). Moreover, those statistical tools often gave candidates whose epigenetic features made them unlikely to have a transcription regulatory function because, even when the tools included epigenetic parameters, these parameters may not have been weighted sufficiently. Our findings also can aid future studies of the incompletely understood roles of genes like *RP11-392O17.1*, *ZBED3-AS1, FAM13A, HOXA11* in adipose tissue and other normal or malignant tissues (26, 56, 69–72). Most importantly, comprehensive biochemically-based prioritization of credible candidates for regulatory SNPs from obesity GWAS can greatly facilitate the demanding experiments required to directly verify allele-specific effects on gene expression linked to inherited obesity risk.

## Materials and Methods

### Obesity GWAS data

Index SNPs associated with obesity, BMI, WHR_adjBMI_, WC_adjBMI_ and HC_adjBMI_ (p < 5 × 10^−8^) were retrieved from 50 studies in the NHGRI-EBI GWAS Catalog (73) (downloaded July 2020; Supplemental Table S1). We expanded the index SNPs to a set of proxy SNPs (*r*^2^ ≥ 0.8, EUR from the 1000 Genome Project Phase 3 (74) by using PLINK v1.9 (75) (https://www.cog-genomics.org/plink/1.9/) and/or LDlink v3.9 (76) (https://ldlink.nci.nih.gov/). For the genes in Table 1, we also extracted imputed SNPs for BMI and WHR_adjBMI_ (20) (p < 5 × 10^−8^) in the 1-Mb region surrounding the genes.

### Epigenomics

Chromatin state segmentation data were from the 18-state Roadmap Epigenome model (8) (https://egg2.wustl.edu/roadmap/data/byFileType/chromhmmSegmentations/ChmmModels/core_K27ac/jointModel/final/. Promoter chromatin refers to strong promoter chromatin (state 1), and enhancer chromatin to strong enhancer chromatin (states 3, 8, 9, or 10). In figures, we visualized these and other data in the UCSC Genome browser (http://genome.UCSC.edu) (13) and slightly modified the color-coding for chromatin state segmentation accordingly to improve clarity. In evaluating promoter vs. enhancer designations, note that these occasionally functionally overlap depending on the cell and DNA context (77). DHS and peaks of H3K27ac were defined as narrowPeaks (Roadmap Project (8), https://egg2.wustl.edu/roadmap/data/byFileType/peaks/consolidatedImputed/narrowPeak/). The 15 non-adipose tissues examined to determine strong promoter/enhancer chromatin and H3K27ac peaks preferentially in SAT vs. ≤ 4 non-adipose tissues are given in Supplementary Table S2. The long-distance chromatin interactions were from GeneHancer tracks (GeneHancer Track Settings (ucsc.edu) (49)) for the LCL or from the Micro-C track at the UCSC Genome Browser (Hi-C and Micro-C Track Settings (ucsc.edu)) for foreskin fibroblasts and embryonic stem cells (40). Adipocytes are mesenchymal stem cells induced to differentiate *in vitro* to adipocytes (8).

### Transcriptomics

Expression ratios were determined from the TPM of SAT or VAT to that of 35 heterologous tissue types (Supplementary Table S3) from the GTEx portal (12). Preferential expression was defined as this ratio for either SAT or VAT > 2 and with a minimum of TPM > 2 for SAT or VAT. Transcript specific expression and single-nuclei expression were also from GTEx (https://gtexportal.org/). Expression data for cell cultures was from strand-specific RNA-seq of total RNA (CSHL long RNA-seq) and 5’ cap analysis of gene expression (http://genome.ucsc.edu (13)). Preadipocytes were primary cultures from SAT.

### Excluding EnhPro SNPs linked to missense SNPs and predicting allele-specific TFBS

The EnhPro SNPs were examined for appreciable linkage to missense SNPs (HaploReg v4.1 (https://broadinstitute.org)) at an LD >0.2 (EUR) to eliminate such SNPs from further consideration as candidate transcription regulatory SNPs. Predictions of allele-specific TFBS from the remaining SNPs were using the TRANSFAC v2020.1 program and database (http://gene-regulation.com) with manual curation as previously described (78). Each TF for a matching TFBS had to have TPM ≥ 2 in SAT or VAT. In addition, we used manual curation to retain only those TRANSFAC TFBS predictions for which all the conserved positions had exact matches to the SNP-containing sequence and for which no more than one base in a partly conserved position had only a partial match (at least 20% as good as the best PWM match). We also looked for evidence from a TF ChIP-seq database (40) for TF binding to oligonucleotide sequences containing our Tier-1 SNPs but no evidence of such binding was found. This is not surprising given the severe limitations on the size of available databases for TF binding in various tissues and cell types.

### Statistical analyses to look for potential causal regulatory SNPs

For the SMR and HEIDI (9), PAINTOR (22), and eCAVIAR (21) analyses, we used the top associated SNPs (p < 5 × 10^−8^) from sex-combined WHR_adjBMI_ GWAS summary data (20). For the SMR and HEIDI analysis, we employed default values of p_eQTL_ < 5 × 10^−8^ for SMR and p_eQTL_ < 1.57 × 10^−3^ for HEIDI, a Bonferroni corrected threshold of p_SMR_ < 2.16 × 10^−6^ and p_HEIDI_ > 0.05. For PAINTOR and eCAVIAR analyses, we only considered significant GWAS SNPs within the 1- to 2-Mb neighborhood of Tier-1 genes in Table 1. For PAINTOR, we integrated overlap of the SNPs with exonic DNA, chromatin segmentation states 1, 3, 8, 9, or 10 (18-state model (8)), or narrow peaks of DHS and H3K27ac in SAT (13). For colocalization eCAVIAR, we included SNPs with significant eQTLs in SAT located within the 1- to 2-MB neighborhood of the genes in Table 1. For COJO in genome-wide complex trait analysis (GCTA) (18, 19), we used the default parameters to evaluate the changes in the association of Tier-1 SNPs with obesity measures by conditioning on any neighboring EnhPro SNPs that lacked a predicted allele-specific TFBS. In addition, we assessed the Tier-1 SNPs for the *HOXC4/C5/C6* subcluster conditional on the *MIR196A2* SNP rs11644913. The reference panel of European individuals (EUR) from the 1000 Genomes Project (phase 3) was used for linkage disequilibrium estimates for all analyses because most GWAS data was for this group.

### Transcription from *RP11-392O17.1* in SAT-MSC

Total RNA was isolated (https://www.foreivd.com/cell-total-rna-isolation-kit-product/) from low-passage MSC derived from SAT that were >95% positive for CD73, CD90, and CD105 and <2% positive for CD45, CD34, CD11b, and CD19 (https://www.cellcook.com). First-strand cDNA synthesis (https://www.uebio.com/productDe_46.html) used ~300 ng of RNA template and included a double-strand DNA-specific DNase in the reaction mixture. PCR (https://www.uebio.com/productDe_443.html) was just for 35 cycles or for 35 cycles followed by a second round of 25 cycles on a 1:10 dilution of the first-round product.

## Supporting information

ZhangX_ObesityRisk_RegSNPs_SupplTables_F_10_18_21

ZhangX_Obesity-Risk_RegSNPs_SupplFigs_F_10_29_21

## Acknowledgements

We thank Dr. Carl Baribault for his help with collecting and editing obesity GWAS data collection and, Ms. Yu Liu and Dr. Shu Ran for their help with TFBS prediction evaluation. This study was partially supported by grants from the National Institutes of Health (P20GM109036, R01AR069055, U19AG055373, and R01MH104680).

