## Supplementary material for "Epigenomic and transcriptomic prioritization of candidate obesity-risk regulatory GWAS SNPs": ZhangX_Obesity-Risk_RegSNPs_SupplFigs_F_10_29_21

This file contains the following figures (Supplementary Tables are in a separate file)

Figure S1. Part of *CABLES1* intron 1 showing two obesity GWAS-derived credible transcription regulatory SNPs (Tier-1 SNPs).

Figure S2. *CABLES1* shows alternative tissue-specific transcription start site usage that may be modulated by upstream and downstream enhancer chromatin containing Tier-1 SNPs.

Figure S3. *PEMT* has two intragenic Tier-1 SNPs, one of which might modulate usage of alternative transcription start sites (TSS).

Figure S4. *PEMT* displays alternative tissue-specific transcription start site usage that may be modulated by enhancer chromatin containing Tier-1 SNPs.

Figure S5. *PC* has a tissue-specific TSS near its Tier-1 SNP whose usage excludes transcription of a miRNA precursor.

Figure S6. Transcription factor binding to a novel cell type-specific TSS near TSS 2 of *PC* and its adjacent Tier-1 SNP.

Figure S7. The six Tier-1 SNPs associated with *FAM13A* are upstream or downstream of TSS 1.

Figure S8. A Tier-1 SNP in the 3' untranslated region of *HOXA11* might help regulate expression of *HOXA10* and *HOX11-AS* as well as *HOXA11* in SAT through enhancer chromatin overlapping this SNP.

Figure S9. SNPs upstream of *RP11-392O17.1* are much closer to this gene than to *LYPLAL1,* a gene with which they have been often associated in the literature.

Figure S10. Evidence for *RP11-392O17.1* expression in a few cell types but not for *LYPLAL1-AS1* expression in any examined cell type

Figure S11. SAT-MSC RT-PCR products from the 5' end of *RP11-392O17.1.*

Figure S12. One of the two *ZBED3-AS1* Tier-1 SNPs is very close to a constitutive CTCF site and the other is near a cell type-specific CTCF site.

Figure S13. Single-nucleus RNA-seq of cell types within various human tissues shows that adipocytes have the most frequent expression of *ZBED3-AS1.*

Figure S14. The formation of a topologically associating domain (TAD) with boundaries near *ZBED3-AS1* Tier-1 SNP rs9293708 and the promoter of *WDR41* was seen in foreskin fibroblasts but not in embryonic stem cells.

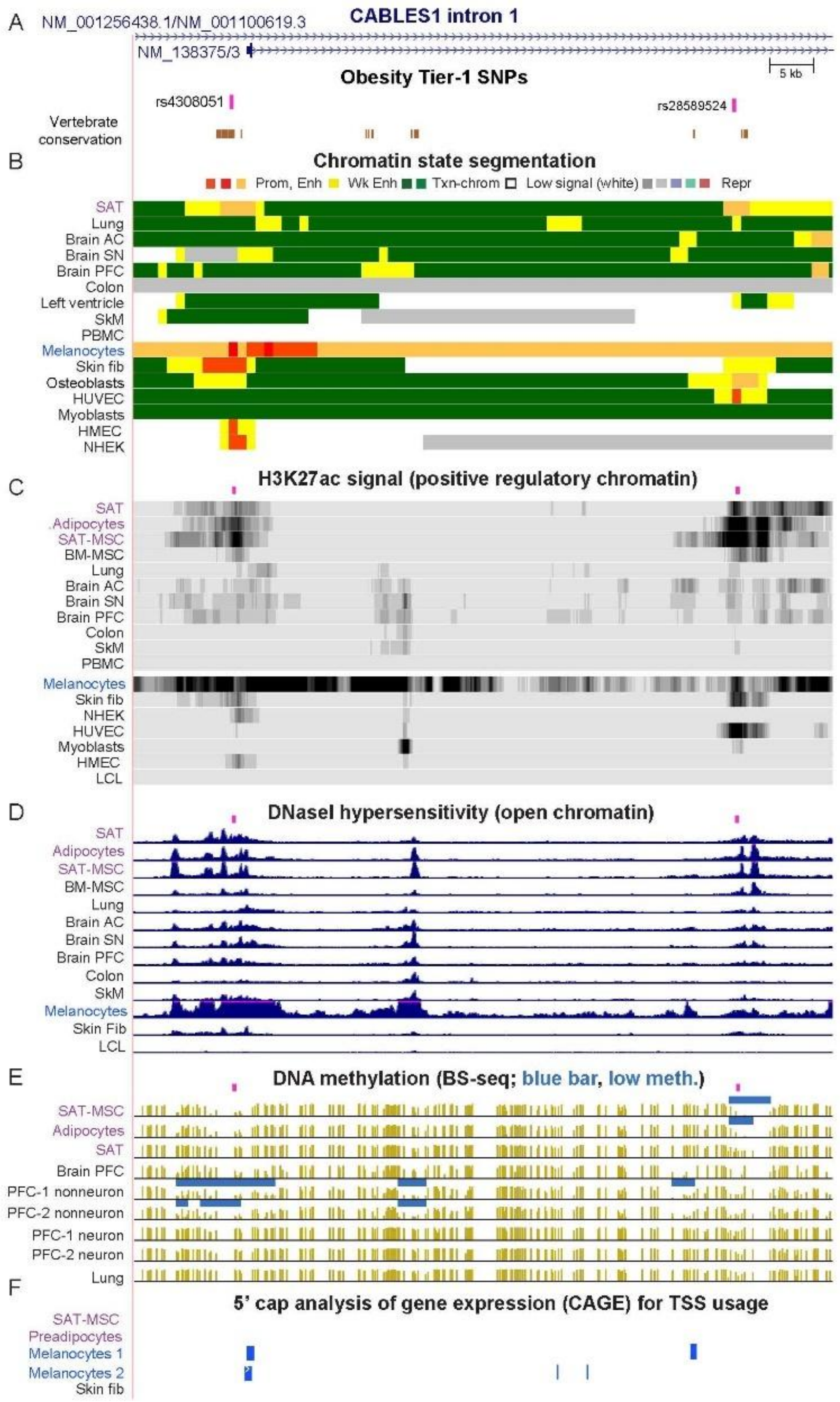


**Figure S1**. **Part of *CABLES1* intron 1 showing two obesity GWAS-derived credible transcription regulatory SNPs (Tier-1 SNPs). A**. A 16-kb region (chr18:20,733,259-20,749,076) containing two of the obesity GWAS-derived Tier-1 SNPs and TSS 2. Element conservation among vertebrates is shown to indicate the overlap of rs4308051 with a conserved DNA sequence. The RefSeq unique identifiers for two coding transcript isoforms are given. **B**. Roadmap-derived chromatin state segmentation: strong promoter (Prom) or enhancer (Enh) chromatin; weak enhancer chromatin (Wk Enh); chromatin with the H3K36me3 mark of actively transcribed regions (Txn-chrom); chromatin with low signals for the assessed histone modifications; repressed (Repr) chromatin. **C**. H3K27ac enrichment profiles. **D**. Profiles of open chromatin (DNaseI hypersensitivity). **E**. DNA methylation profiles from whole-genome bisulfite sequencing; blue horizontal bars indicate regions of significantly low methylation relative to the rest of the same genome. **F**. Transcription start site profiling by 5' cap analysis gene expression (CAGE, TSS HMM clusters (1)) from the UCSC Genome Browser showing much transcription initiation from TSS 2 in melanocytes. There were also high levels of transcription initiation from TSS 1 and overall levels of RNA were much higher in melanocytes than in any other cell type in the ENCODE database at the UCSC Genome Browser (http://genome.ucsc.edu, data not shown). SAT, subcutaneous adipose tissue. AC, anterior cortex; SN, substantia nigra; PFC, prefrontal cortex; SkM, skeletal muscle; PBMC, peripheral blood mononuclear cells; Fib, fibroblasts; HUVEC, umbilical cord endothelial cells; HMEC, mammary epithelial cells; NHEK, epidermal keratinocytes; SAT-MSC, SAT-derived mesenchymal stem/stromal cells; BM-MSC, bone marrow-derived MSC; LCL, lymphoblastoid cell line (GM12878); PFC-1 or -2 nonneuron, biological duplicates of cells derived from brain PFC depleted of neurons; PFC-1 or -2 neuron, biological duplicates of cells derived from brain PFC enriched for neurons. All tracks were from the UCSC Genome browser with hg19 coordinates.

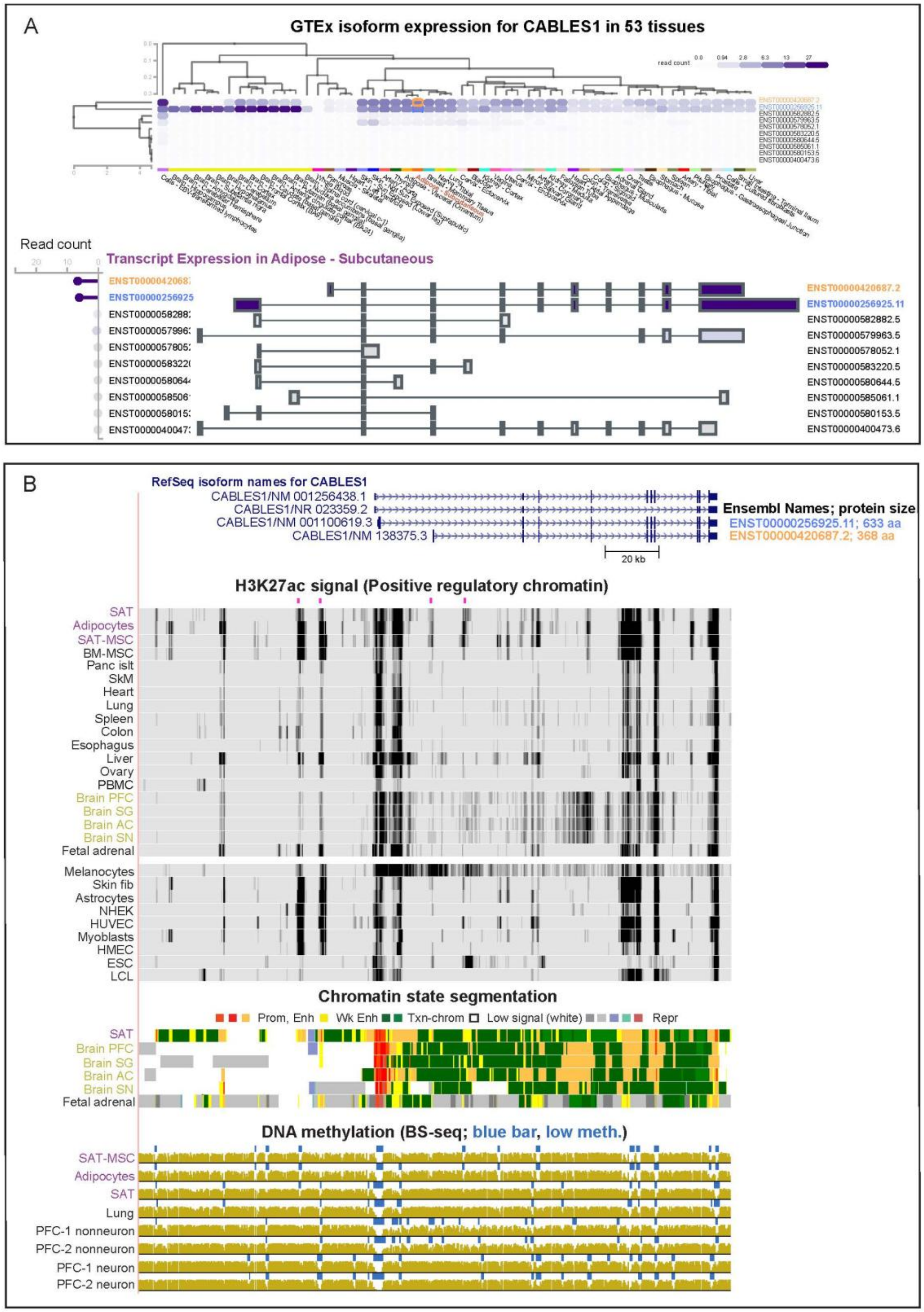


**Figure S2**. ***CABLES1* shows alternative tissue-specific transcription start site usage that may be modulated by upstream and downstream enhancer chromatin containing Tier-1 SNPs. A**. Isoform expression for *CABLES1* from the GTEx database (https://www.gtexportal.org/ V8). The Ensembl transcript names for the main isoforms seen in SAT are indicated by orange (starting at TSS 2) or blue font (starting at TSS 1), and their diagrammed read counts in SAT are circled in orange or blue. **B**. The 218-kb region containing *CABLES1* and its four Tier-1 SNPs (chr18:20,627,684-20,845,426; rs4445996, rs9966093, rs4308051, and rs28589524,). The four gene isoforms are shown as well as H3K27ac signal, chromatin state segmentation and whole-genome bisulfite sequencing as in Figure S1. Short pink bars indicate the location of Tier-1 SNPs. The sizes of the corresponding proteins in amino acids (aa) are indicated at the right. Note brain-specific H3K27ac and chromatin state profiles. Panc islt, pancreatic islets; ESC, embryonic stem cells.

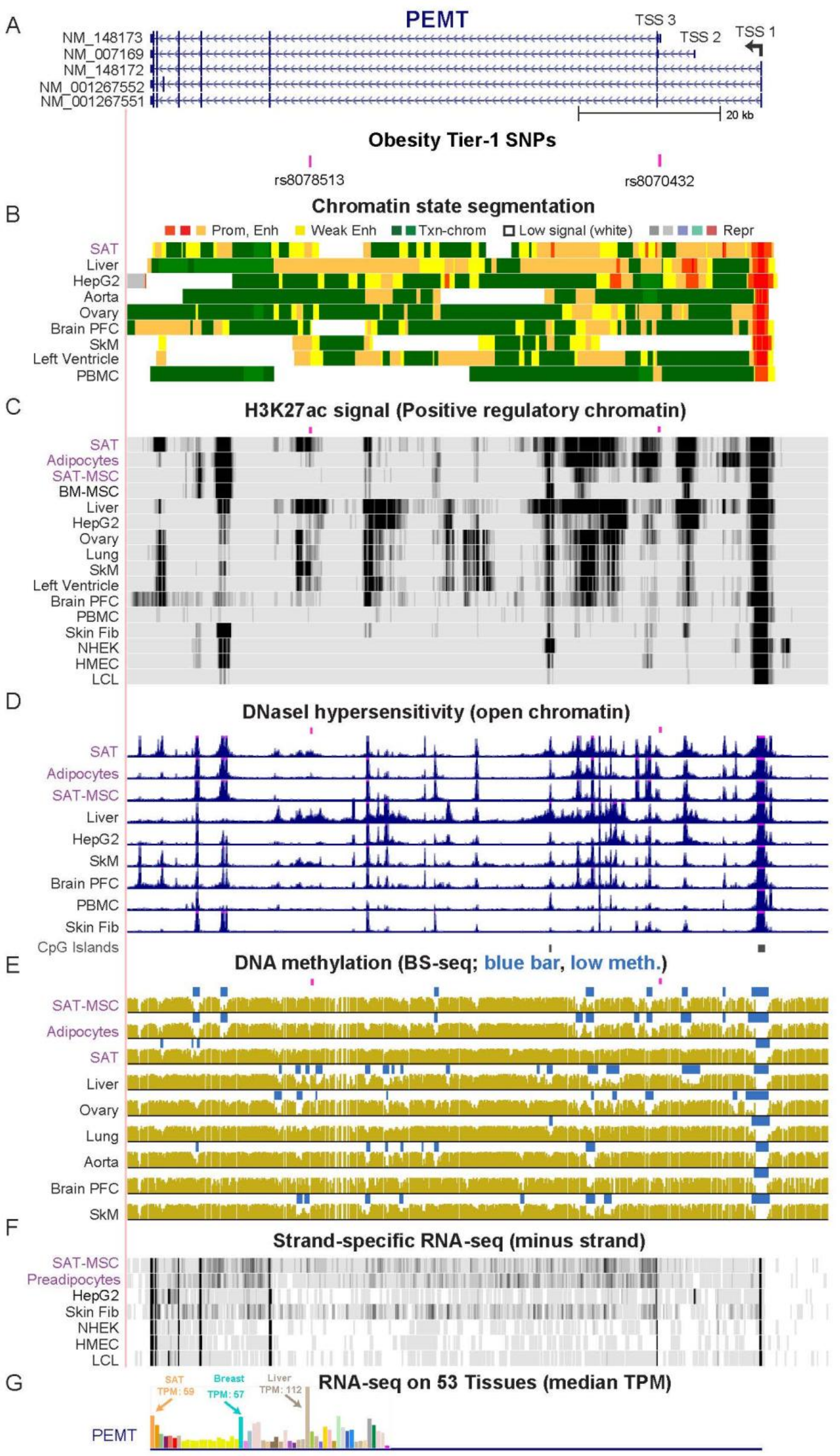


**Figure S3**. ***PEMT*** **has two intragenic Tier-1 SNPs, one of which might modulate usage of alternative transcription start sites (TSS). A.** The 99-kb region (chr17:17,405,553-17,504,445) containing five gene isoforms of *PEMT* that originate from three different transcription start sites (TSS 1, 2, and 3) as well at the two Tier-1 SNPs for this gene (rs8078513 and rs8070432). **B** - **F**. Chromatin state segmentation, H3K27ac, DNaseI hypersensitivity, bisulfite-seq as in Figure S1 with the addition of strand-specific RNA-seq profiles (2). Short pink bars above tracks in panels **C**, **D**, and **E**, positions of Tier-1 SNPs. **G**. Tissue RNA-seq shown as a bar graph with median TPM from hundreds of biological replicates (first two orange bars are SAT and visceral adipose tissue, VAT; all yellow bars are brain samples; see Supplementary Table S3 for detailed data on 34 of these tissues). HepG2, hepatocellular carcinoma cell line.

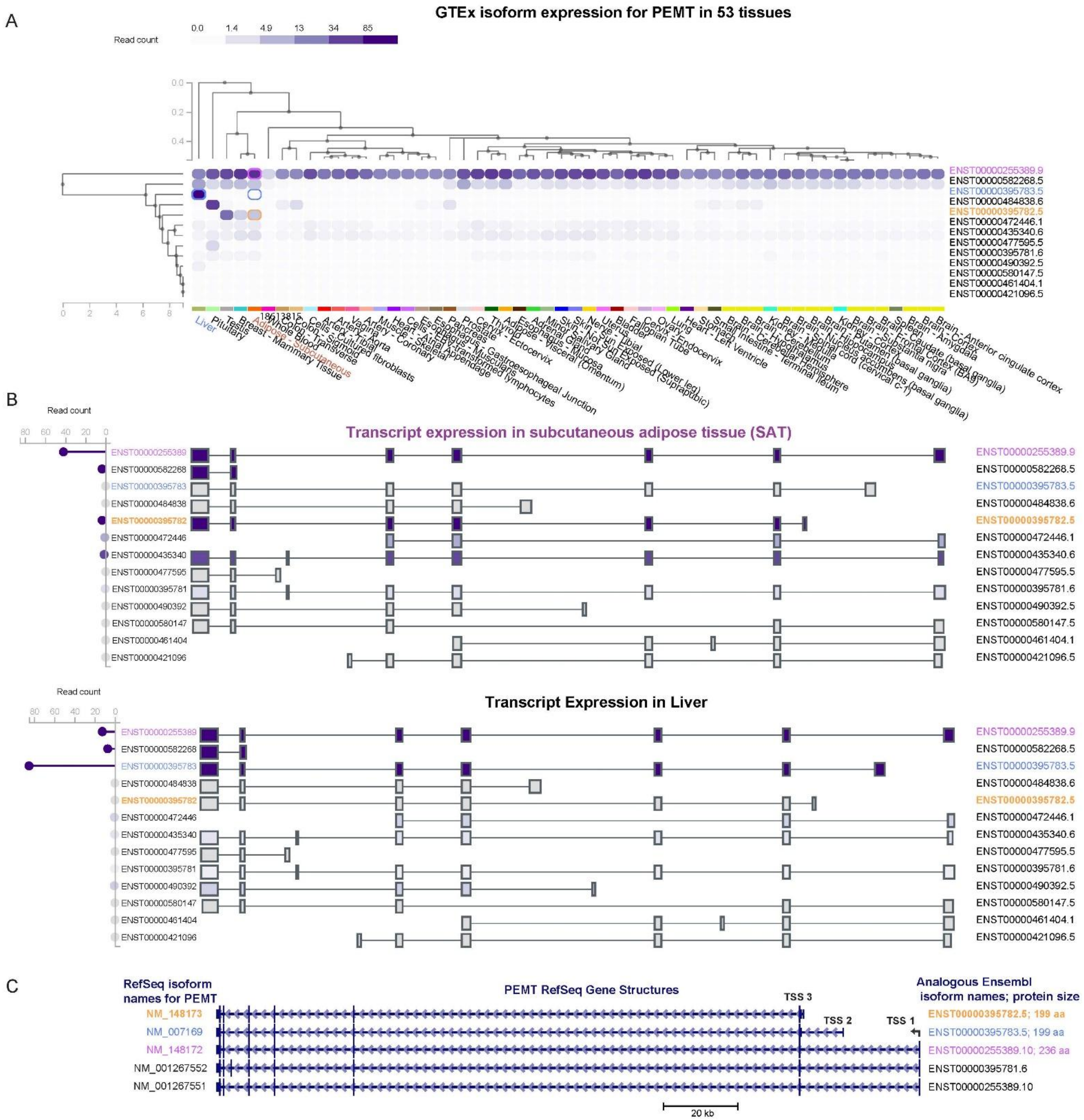


**Figure S4**. ***PEMT* displays alternative tissue-specific transcription start site usage that may be modulated by enhancer chromatin containing Tier-1 SNPs. A**. GTEx data for expression of different isoforms in many different tissues as in Figure S2; the TSS is on the right. **B**. More details for SAT and liver. RNA isoforms displaying different relative frequencies in SAT vs. liver are indicated by colored font with the corresponding colors used to circle the diagrammed count data for SAT and liver in panel **A**. **C**. The sizes of the encoded proteins are indicated on the right of this panel.

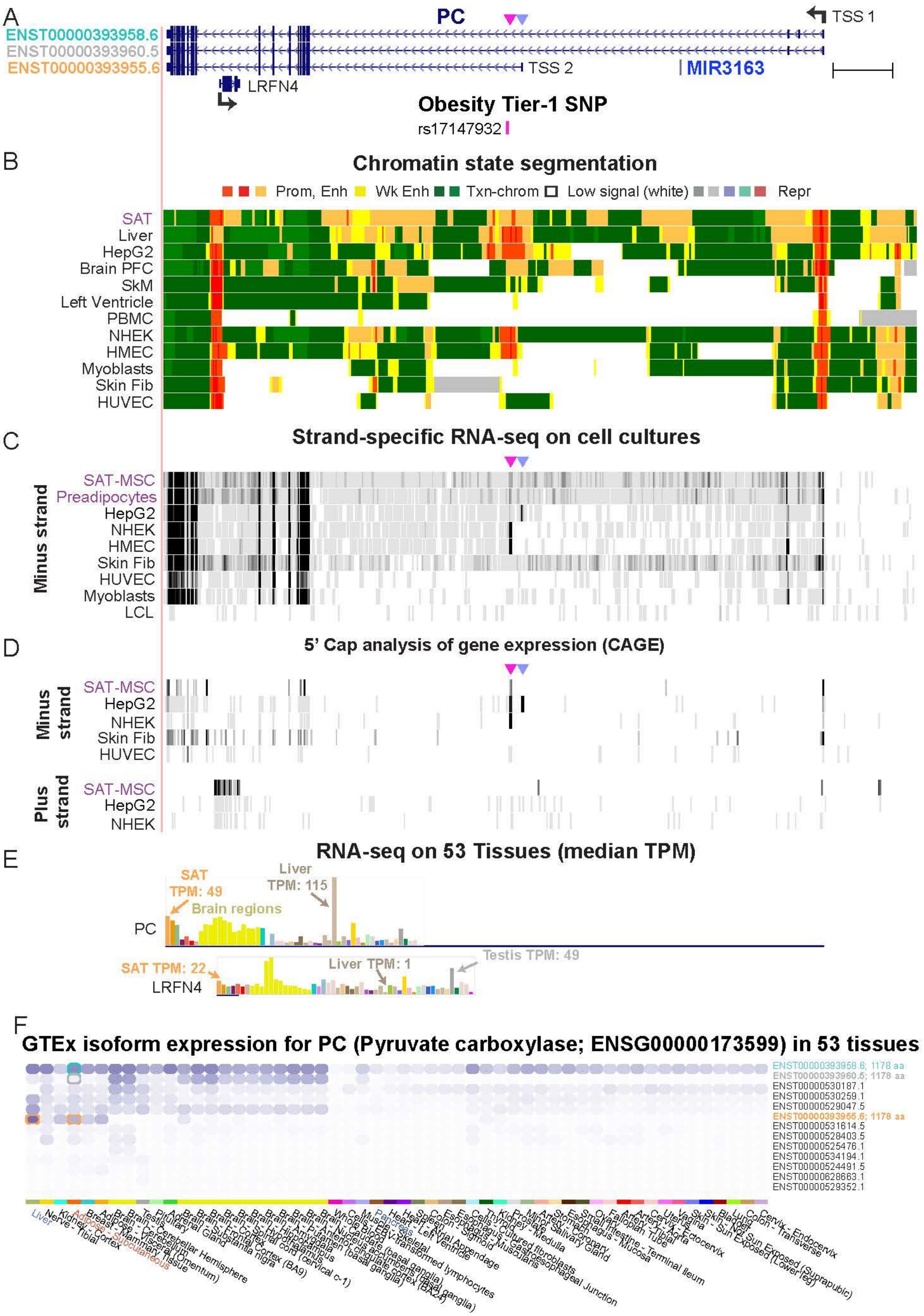


**Figure S5.** ***PC* has a tissue-specific TSS near its Tier-1 SNP whose usage excludes transcription of a miRNA precursor**. **A**. A 126-kb region containing *PC* (*Pyruvate Carboxylase*, *ENSG00000173599*; chr11:66,615,239-66,741,500), its one Tier-1 SNP, two TSS, one intragenic miRNA-encoding gene (*MIR3163* in the minus strand), and another small intronic gene, which is antisense to *PC*, *LRFN4*). Pink triangle, a novel TSS (referred Figure S6 as TSS 3), 0.7 kb upstream of the Tier-1 SNP rs17147932; blue triangle, the previously documented proximal TSS, TSS 2 of *PC* (see Supplementary Figure S6). **B**, **C**, and **D**. Chromatin state segmentation, H3K27ac, RNA-seq, and CAGE profiles as in Figure S1, except that the CAGE signal is shown. **E**. Bar graphs with tissue RNA-seq data for *PC* and its intragenic *LRFN4* as in Figure S3. **F**. RNA-seq (GTEx) showing tissue-specific isoform expression (compare SAT and liver) involving the use of alternate TSS. In addition, the sizes of the encoded proteins are given on the right.

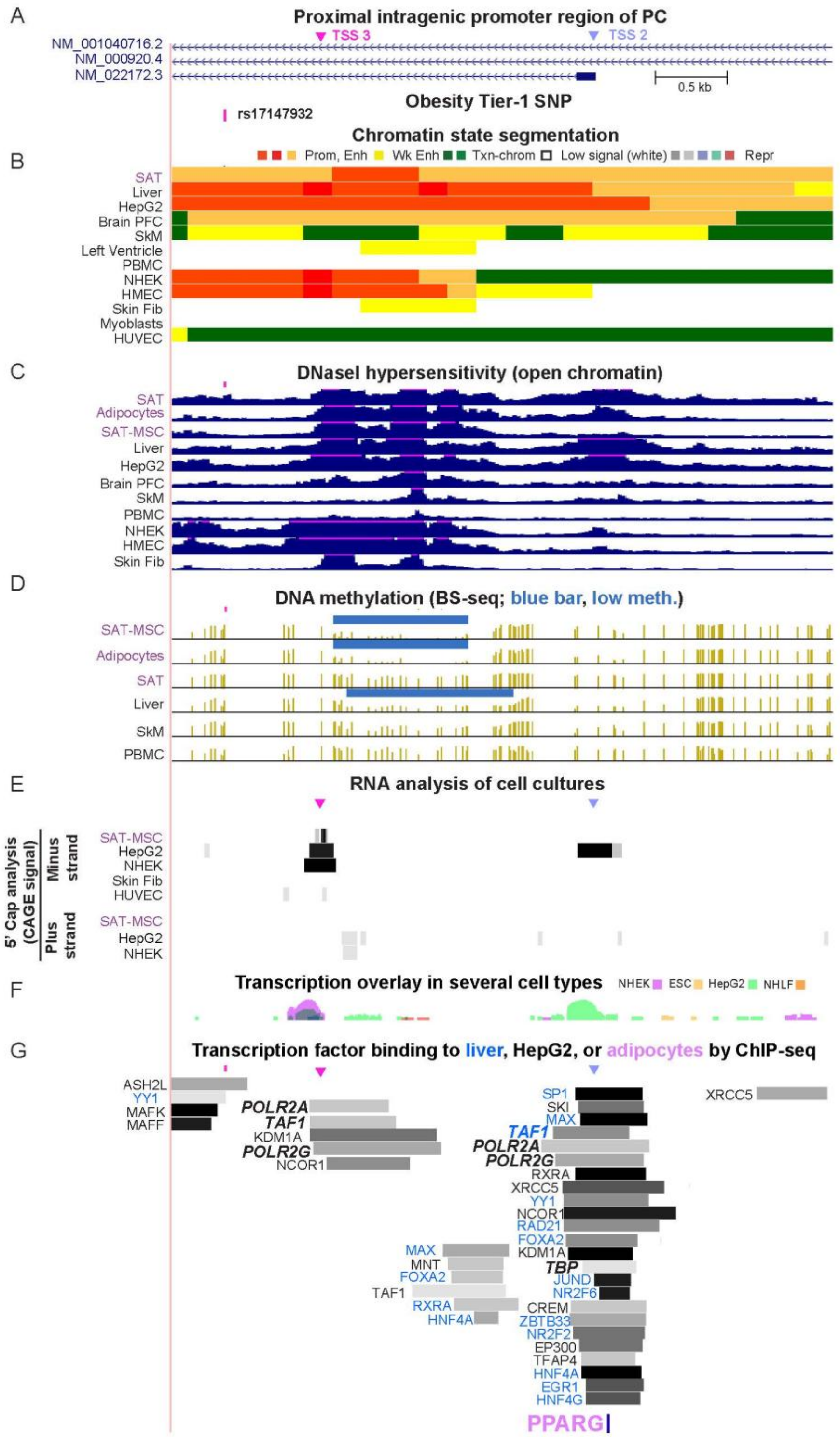


**Figure S6**. **Transcription factor binding to a novel cell type-specific TSS near TSS 2 of *PC* and its adjacent Tier-1 SNP. A**. A 4.6-kb region (chr11:66,672,491-66,677,062) around the second previously documented TSS of PC; **B** - **F**. Chromatin state segmentation, DNaseI hypersensitivity, bisulfite-seq, and cell culture CAGE profiles for the indicated strand as in Figures S1 and S5. The horizontal pink lines on top of DHS peaks in **C** indicate off-scale peaks. **F**. Another cell culture RNA-seq database at the UCSC Genome Browser (ENCODE Regulation Transcription Track Settings (ucsc.edu)), which shows the signal from four cell cultures superimposed on one another according to the indicated color code. The combined data in Panels **E** and **F** indicate that the previously undescribed TSS 3 was used frequently in these cell cultures. **G**. Transcription factor (TF) binding from chromatin immunoprecipitation/next-gen sequencing (ChIP-seq) to this region for liver and HepG2 cells (ENCODE Regulation Txn Factr ChIP E3 Track Settings (ucsc.edu)). Also shown is the ChIP-seq binding of the adipogenesis-associated TF PPARG in adipocytes generated *in vitro* from SAT-MSC (UniBind Track Settings (ucsc.edu) (3)). For ease of viewing, most, but not all TF binding sites from ChIP-seq are shown. Color coding indicates the sample type for the observed TF binding; the intensity of the gray/black signal denotes the strength of the signal. The names of the RNA polymerase subunits or basal transcription initiation factors that support transcription initiation at TSS 2 or the novel TSS 3 are shown in bold italics. Pink and blue triangles in **A**, **E**, and **G**, TSS 3 and TSS 2, respectively, which are 0.7 and 2.5 kb upstream of the Tier-1 SNP rs17147932.

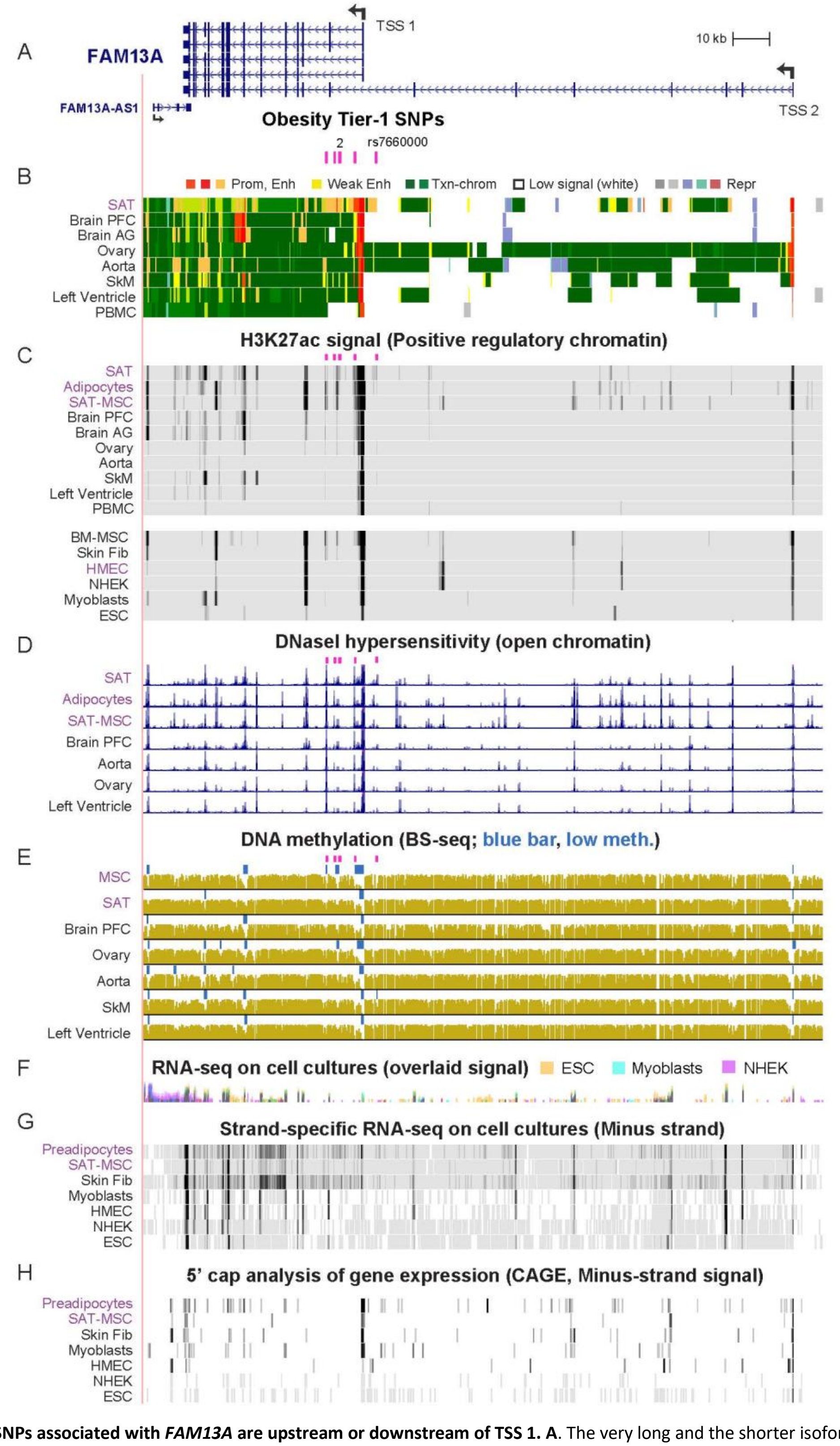


**Figure S7**. **The six Tier-1 SNPs associated with *FAM13A* are upstream or downstream of TSS 1. A**. The very long and the shorter isoforms of *FAM13A* in a 369-kb region (chr4:89,625,479-89,994,224) containing six Tier-1 SNPs (rs10012045, rs2869949, rs11097198, rs6532080, rs13133548, and rs7660000); the SNP not included in Figure 3 is identified and the two SNPs closely adjacent to one another are indicated. **B** - **H**. Chromatin state segmentation, H3K27ac, DNaseI hypersensitivity, bisulfite-seq, cell culture RNA-seq shown by color-coded overlaid signals, strand-specific cell culture RNA-seq, and CAGE profiles for the minus strand as in previous figures. The CAGE profiles for cell cultures show cell type-specific differential use of TSS 1 and TSS 2 consistent with RNA-seq profiles for these cells. There were no CpG islands in this region. Note the different relative sizes of the promoter chromatin regions among some tissues, e.g., aorta vs. SAT, which reflects their relative frequency of TSS usage as seen in the GTEx database (not shown; https://gtexportal.org/home/). The vertical viewing scale for DNaseI hypersensitive peaks here is 0 - 20 instead of 0 - 7 in Figure 3.

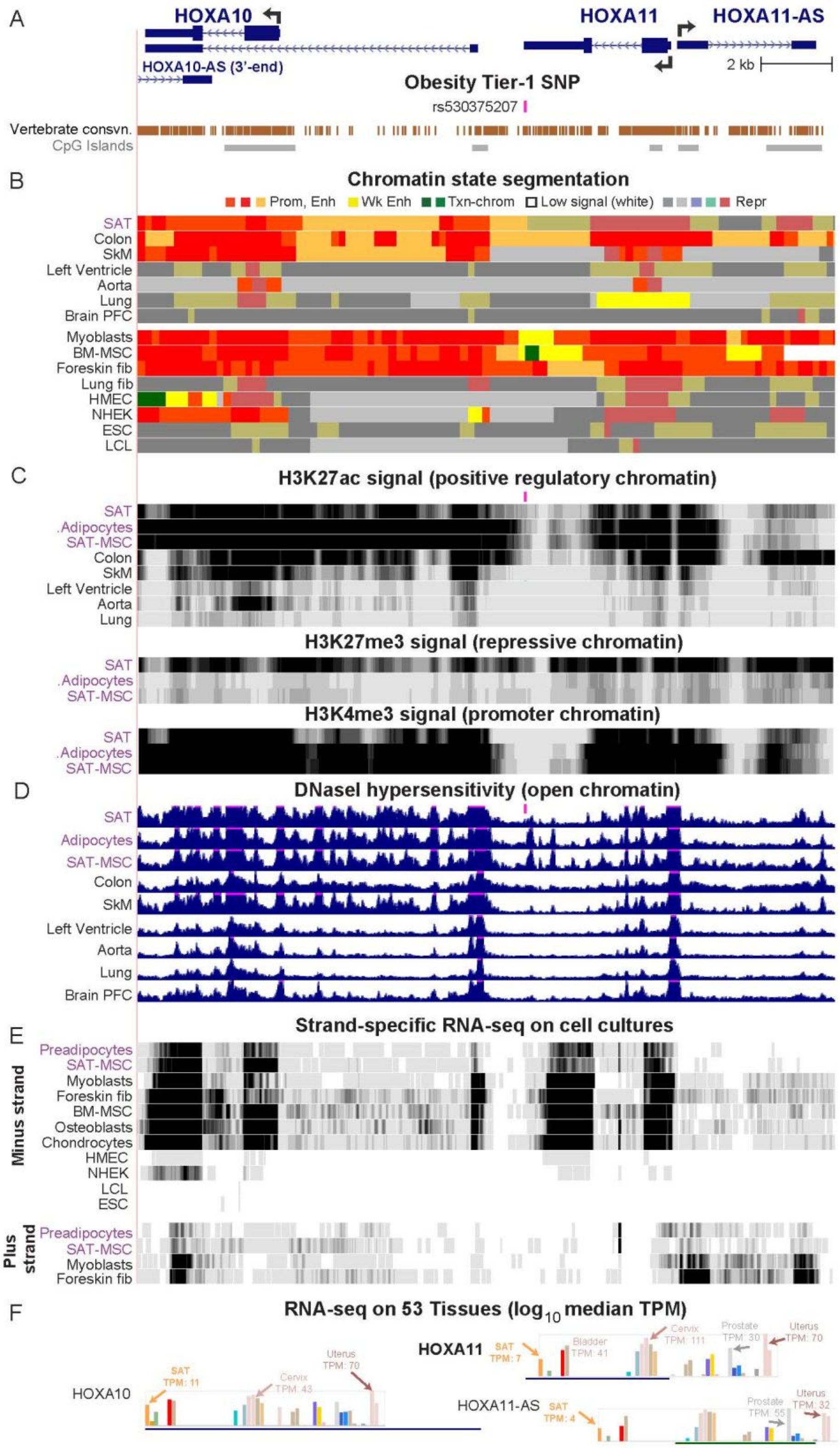


**Figure S8**. **A Tier-1 SNP in the 3' untranslated region of *HOXA11* might help regulate expression of *HOXA10* and *HOX11-AS* as well as *HOXA11* in SAT through enhancer chromatin overlapping this SNP. A**. A 19-kb region (chr7:27,210,012-27,229,330) from the *HOXA* gene cluster containing Tier-1 SNP (rs530375207) showing vertebrate element conservation and CpG islands. **B** - **F**. Chromatin state segmentation, H3K27ac, DNaseI hypersensitivity, strand-specific cell culture RNA-seq, and GTEx bar graphs are as in Figures S3 and S5 except that unlike the other GTEx bar graphs which depict the TPM in linear scale, panel **F** shows it in $log_{10}$ scale to better compare the tissue-specificity of the three genes. In addition, panel C also shows profiles for H3K27me3 (repression) and H3K4me3 (promoter chromatin when there is also H3K27ac enrichment). The H3K27me3 profiles indicate that although the bidirectional promoter region of *HOXA11* and *HOXA11-AS* in SAT appears to be bivalent/repressed promoter chromatin (gray-red color), the histone modification signal from their promoter region is likely to be due to a mixture of cells with active promoter chromatin and other cells with repressed chromatin in this region. This would explain the considerable RNA-seq signal from SAT for these two genes.

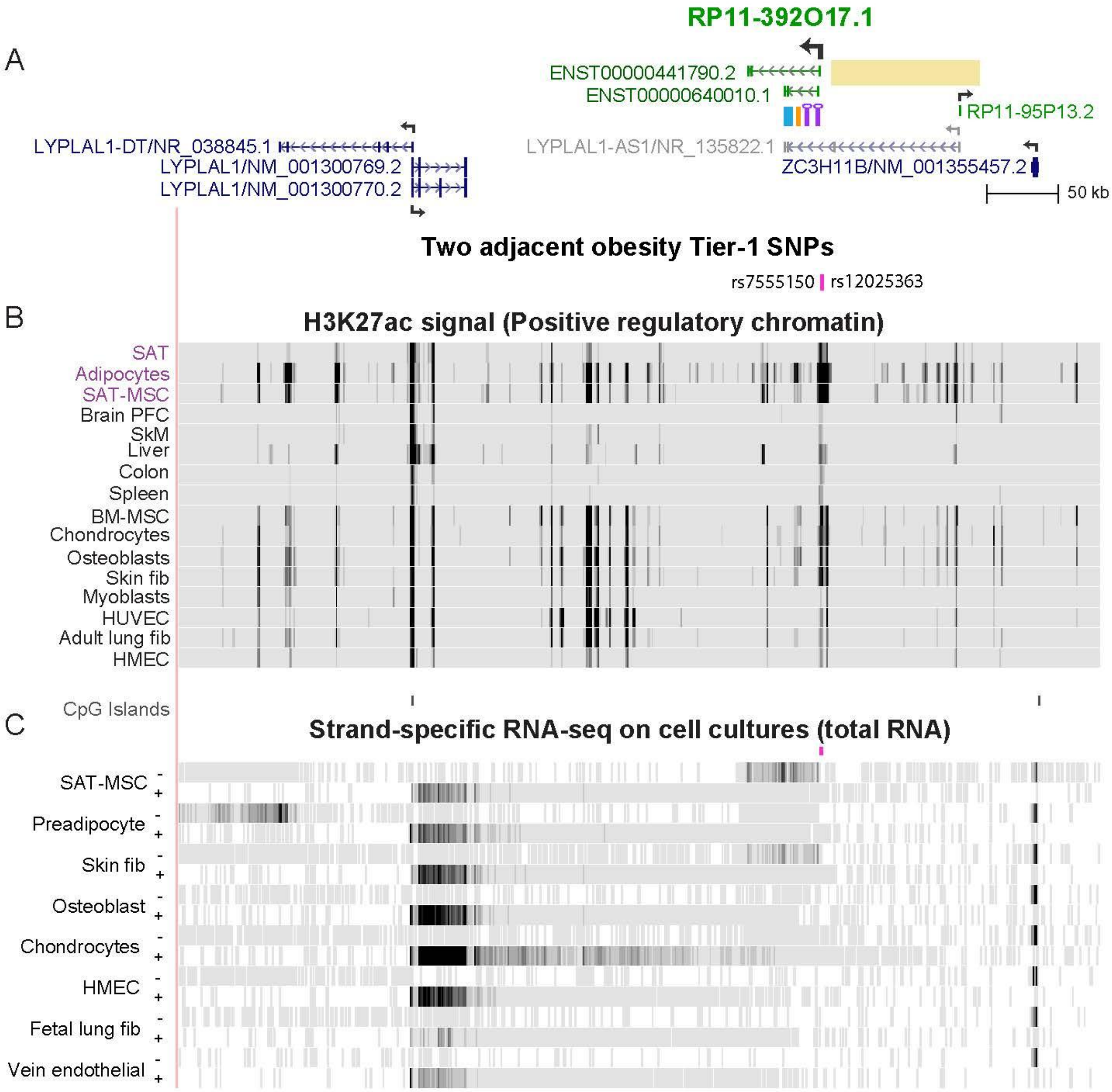


**Figure S9**. **SNPs upstream of *RP11-392O17.1* are much closer to this gene than to *LYPLAL1,* a gene with which they have been often associated in the literature**. **A**. The 0.6-Mb region (chr1:219,183,979-219,829,670) shown contains *RP11-392O17.1*, a gene that is not in the RefSeq database but is in the Ensembl (V104) database as is *RP11-95P13.2* (both shown in green). The RefSeq genes are shown in blue, except for *LYPLAL1-AS1,* which is colored gray because no transcript corresponding to the RefSeq gene labeled *LYPLAL1-AS1* was seen in any of the 16 examined cell types in the ENCODE database even at the highest sensitivity settings (http://genome.ucsc.edu). In many GWAS (4-16), candidate regulatory SNPs located 11 – 118 kb upstream of *RP11-392O17.1* (the region denoted by the tan rectangle) were previously designated as associated with *LYPLAL1* or *LYPLAL1-AS1.* Moreover, *LYPLAL1-AS1* is sometimes used in the literature to denote *RP11-392O17.1* or *LYPLAL1-DT* (another lncRNA gene). For example, Yang *et al.* (17) referred to a 0.5-kb region from *RP11-392O17.1* that they cloned (blue bar in panel **A**) as *LYPLAL1-AS1. RP11-95P13.2*, unlike *RP11-392O17.1*, has barely detectable RNA from SAT, VAT, and breast, negligible RNA in other tissues in the GTEx database (https://www.gtexportal.org/), and undetectable transcription in any cell type even using a high sensitivity setting. The purple lollipops in panel **A** show the approximate positions of *RP11-392O17.1* amplicons used in Figure S10. The adjacent orange bar indicates the position of the amplicon used by Zheng et al. (18) in their study of the gene they call *LYPLAL1-2*, which overlaps *RP11-392O17.1.* They cloned “full-length *LYPLAL1-2*” but did not specify its coordinates, sequence, or size. *LYPLAL1* also does not exhibit any preferential transcription in adipose vs. other tissues, unlike *RP11-392O17.1* (Supplementary Table S10 and Figure S10). **B** and **C**. H3K27ac and strand-specific RNA-seq from total RNA as in previous figures except that vertical viewing range was 0 – 100 instead of the more sensitive 0 – 50 used in Figure 4. The RNA-seq signal seen specifically in preadipocytes and even stronger in myoblasts (not shown) overlaps the last two exons of *LYPLAL1-DT* but extends further and is of unknown function.

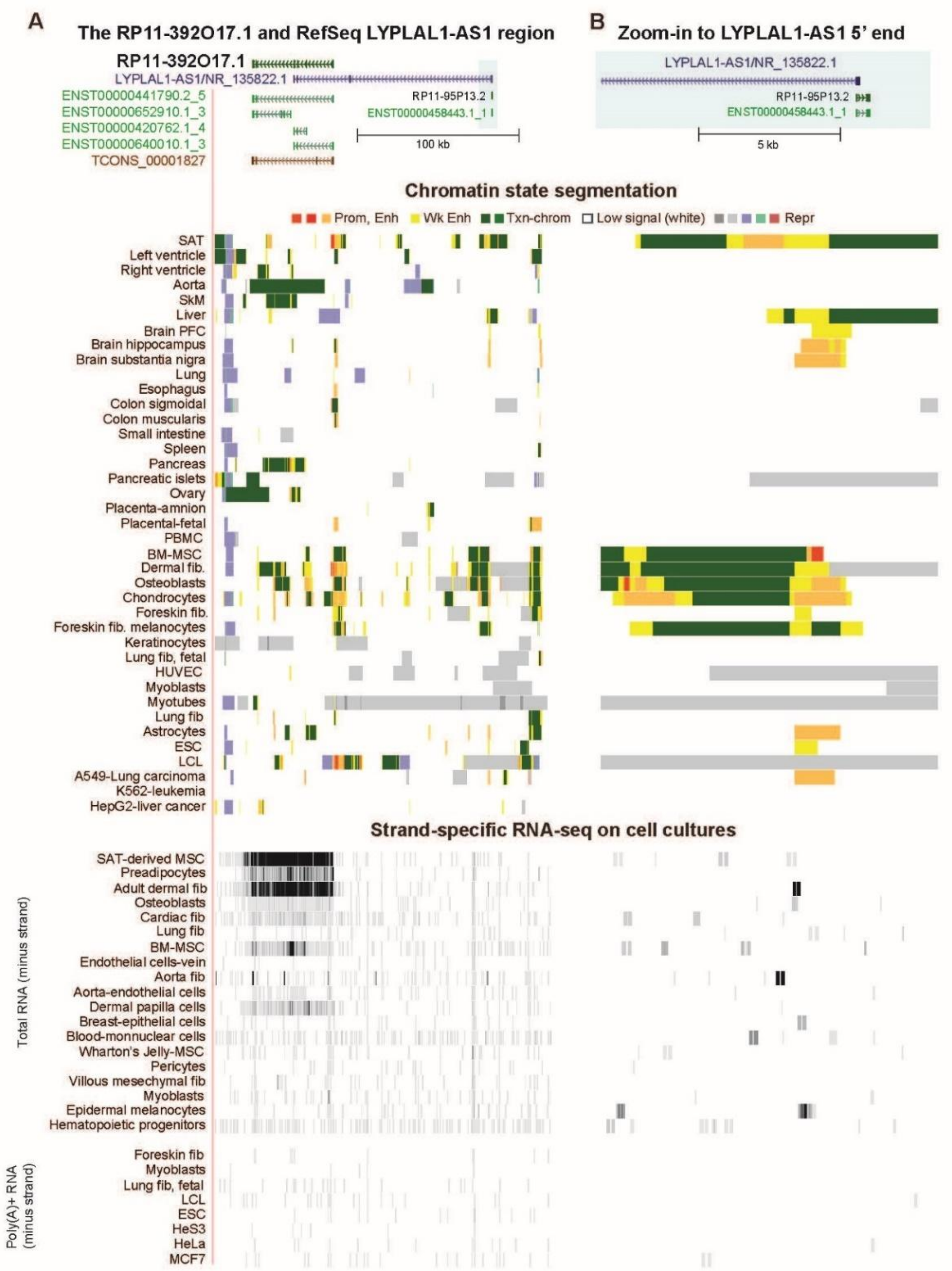


**Figure S10. Evidence for *RP11-392O17.1* expression in a few cell types but not for *LYPLAL1-AS1* expression in any examined cell type. A.** The 280-kb region containing the *RP11-392O17.1* and the RefSeq structure called *LYPLAL1-AS1*; Ensembl isoform structures shown in green; the TCONS gene structure comes from the Human Body Map lncRNAs at the UCSC Genome Browser lincRNAs lincRNA Transcripts Track Settings (ucsc.edu); no TCONS gene structure was found for *LYPLAL1-AS1*. Chromatin state segmentation is as in previous figures. RNA-seq tracks show only the minus strand for both total RNA and poly(A)$^+$ RNA samples with a vertical viewing range of 0 – 30. **B**. Same as panel A but for the 12-kb region (chr1:219,721,578-219,733,398) at the 5' end of *LYPLAL1-AS1* that is highlighted in green in panel A. In addition, the vertical viewing range for the RNA-seq tracks was 0 – 10 to maximally increase sensitivity. Note that all the gene structures shown are minus-strand genes except *RP11-95P13.2*, whose expression was not detectable in plus-strand RNA-seq on these cell cultures (not shown). The light red segment downstream of the *LYPLAL1-AS1* 5' end in the chromatin state track for BM-MSC is state 3 indicating either enhancer or promoter chromatin unlike the strong promoter chromatin (state 1, bright red) immediately upstream of the 5' end of *RP11-392O17.1* in SAT.

A

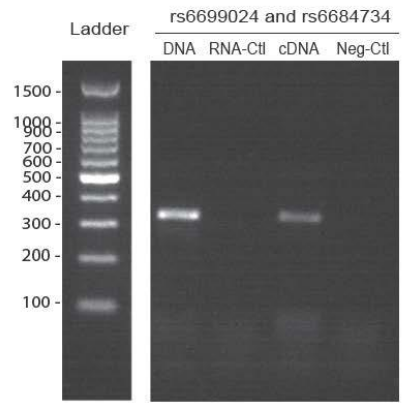


C

Primers for the *RP11-392O17.1* 329-bp amplicon containing SNPs rs6699024 and rs6684734 (5' to 3')

F: CACCATCTTGCCCTCTAACAC

R: CTGGGCTGTGTTTGGTTGTT

B

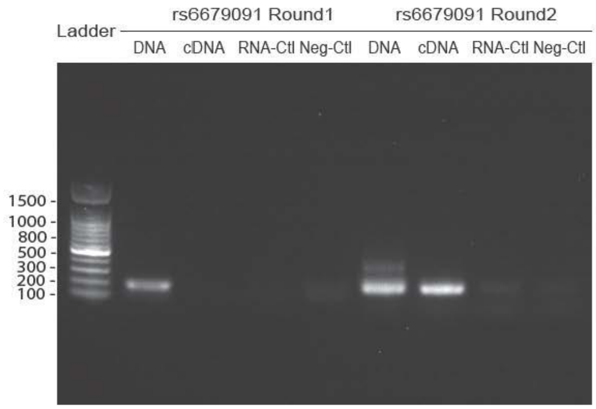


D

Primers for the *RP11-392O17.1* 170- and 152-bp amplicons containing SNP rs6679091 (5' to 3')

First round of PCR

F: GATTCCAACGCGGCTGTC

R: ATGCCCCAGATGAAGCCTAC

Second round of PCR

F: AGAGGCTTCTGGGGCTACAC

R: ATGCCCCAGATGAAGCCTAC

E

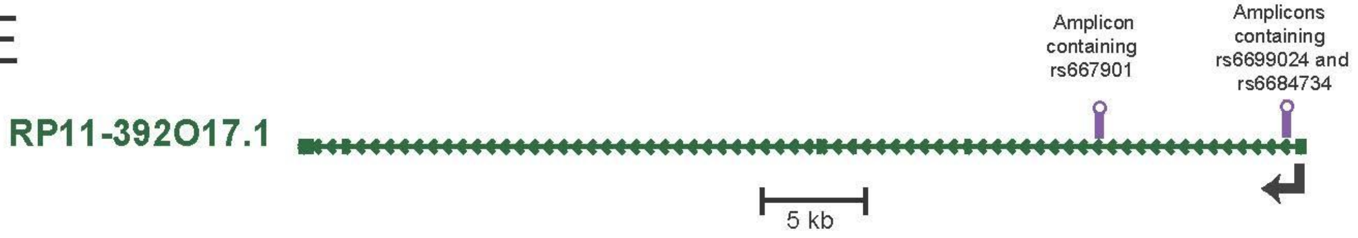


**Figure S11. SAT-MSC RT-PCR products from the 5' end of *RP11-392O17.1*.** **A**. A 329-bp RT-PCR product (chr1:219,631,527-219,631,855) in the 5' end of *RP11-392O17.1* from DNA as a positive control, total RNA as a negative control, cDNA made from total RNA extracted from primary SAT-derived mesenchymal stem/stromal cells, and a no-template negative control. PCR was for 35 cycles. **B**. The second amplicon, a 152-bp RT-PCR product (chr1:219,622,409-219,622,560) obtained after two rounds of PCR (35 cycles and 25 cycles) using the same templates are for panel **A**. **C** and **D**. The primers used for PCR of the cDNA for panels **A** and **B,** respectively. **E.** the structure of the full-length *RP11-392O17.1* gene with the position of the amplicons for RT-PCR indicated. The full-length transcript *ENST00000441790* is predicted to be 1,087 nt but this sequence is labeled as TSL5 in the Ensembl database, which denotes poor support for the positions of its exons. Because the first exon of this transcript consists mostly of Alu repeat sequences, it was not possible to find good PCR primers for it. The indicated SNPs in the amplicons are in high LD with the Tier-1 SNPs but the haplotypes of both RT-PCR products could not be accurately quantitated for allele-specific expression analysis upon Sanger sequencing due to a high background of heterologous sequences.

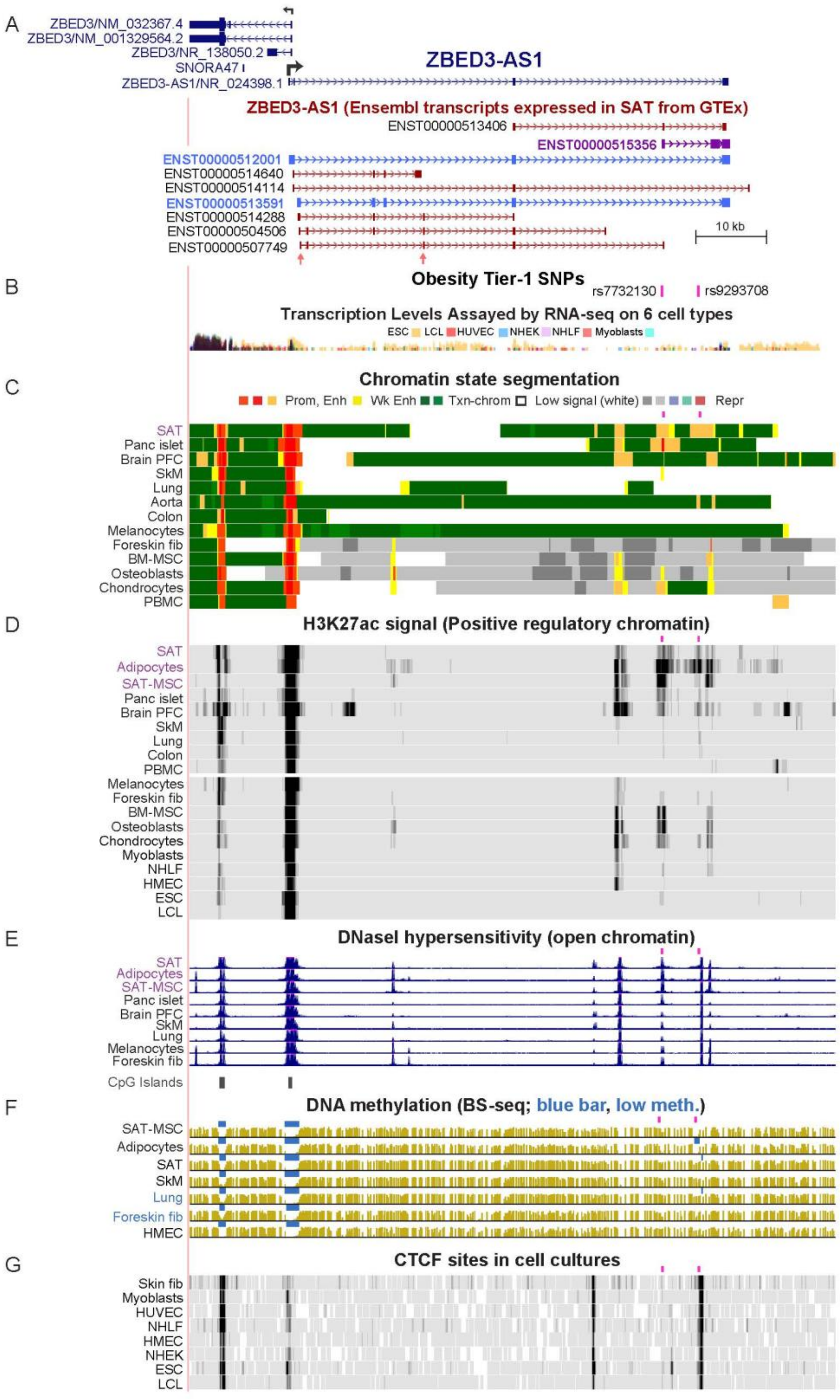


**Figure S12. One of the two *ZBED3-AS1* Tier-1 SNPs is very close to a constitutive CTCF site and the other is near a cell type-specific CTCF site. A.** The 90-kb region containing *ZBED3-AS1* and *ZBED*3 (chr5:76,368,810-76,459,249) showing all their RefSeq isoforms, the Ensembl transcripts for *ZBED3-AS1,* and the two obesity Tier-1 SNPs, rs7732130 and rs9293708. The Ensembl transcripts that are in light blue are the most highly transcribed ones in SAT according to the GTEx database (https://www.gtexportal.org/). The vertical pink arrows under the last Ensembl transcript are the positions of the RT-PCR primers used by Xu *et al*. (19) in their study of *ZBED3-AS1* (which they called *lnc13728*) in differentiating SAT-MSC. **B**. RNA-seq (not strand-specific) profiles from six cell cultures (embryonic stem cells, a lymphoblastoid cell line, umbilical vein endothelial cells, epidermal keratinocytes, lung fibroblasts, and myoblasts) given as a color-coded overlay. This profile shows the high expression of *ZBED3* in all these cell types compared with their low expression of *ZBED3-AS1*. **C** – **F**. Chromatin state segmentation, H3K27ac, DNaseI hypersensitivity, bisulfite-seq, as in Figure S1. **G**. CTCF binding by ChIP-seq (ENCODE project in the UCSC Genome Browser).

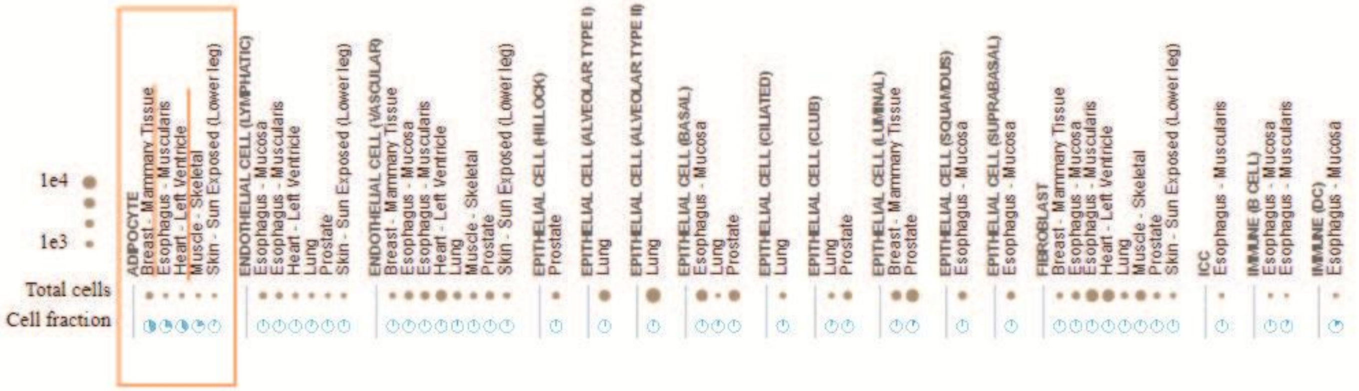


**Figure S13. Single-nucleus RNA-seq of cell types within various human tissues shows that adipocytes have the most frequent expression of *ZBED3-AS1*.** Data are from the GTEx database (https://www.gtexportal.org/) single-nucleus RNA-seq analysis from the indicated tissue. The orange box highlights the data for adipocytes.

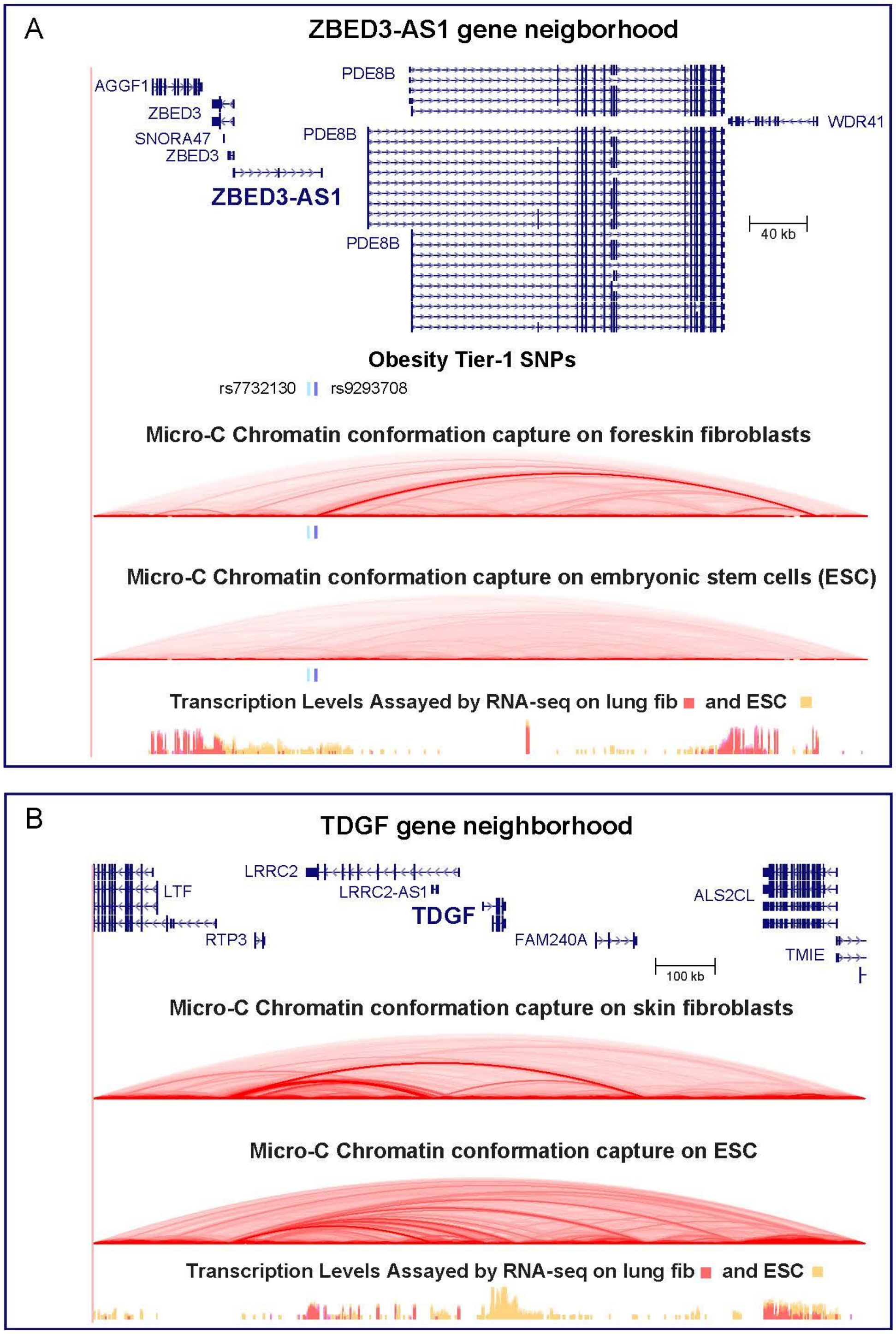


**Figure S14. The formation of a topologically associating domain (TAD) with boundaries near *ZBED3-AS1* Tier-1 SNP rs9293708 and the promoter of *WDR41* was seen in foreskin fibroblasts but not in embryonic stem cells. A**. Micro-C chromatin looping from the UCSC Genome Browser (http://genome.UCSC.edu) seen in the 0.5-Mb *ZBED3-AS1* gene neighborhood; these tracks were visualized in hg38, chr5:76,989,903-77,528,096; in Figure 5, they were transposed to hg19 to combine them with other tracks seen only in hg19. At the bottom are non-strand specific RNA-seq shown in color-coded overlay for lung fibroblasts (skin fibroblasts not available) and H1 embryonic stem cells (ESC). All the RefSeq isoforms for *PDE8B* are shown. Light blue and dark blue bars, the positions of Tier-1 SNPs rs7732130 and rs9293708, respectively. **B**. Micro-C and RNA-seq as in panel A but for the *TDGF* gene neighborhood to serve as a positive control for the detection of ESC-specific chromatin looping. The RNA-seq tracks show the ESC-specific expression of *TDGF*.
